# Multimodal gene and targeted drug therapy for chronic myelogenous leukemia: Computational target analysis and therapeutic validation

**DOI:** 10.1101/2025.05.26.656072

**Authors:** Margaret L. Lugin, Winnie Lei, Rebecca T. Lee, Jee Young Chung, Nicholas M. Katritsis, Woochang Hwang, Angela G. Fleischman, Namshik Han, Young Jik Kwon

## Abstract

Developing an efficient and safe therapy necessitates a mechanistic understanding of the complex underlying pathology and manipulation of the multiple pathways at the molecular and genetic level. Network-based simulation of chronic myeloid leukemia (CML), a relatively well-understood cancer model, revealed the dynamics of simultaneously expressing pro-apoptotic BIM and silencing pro-survival MCL-1 in combination with the BCR-ABL-targeted tyrosine kinase inhibitor dasatinib. Viral/nonviral chimeric nanoparticles (ChNPs) composed of a BIM-expressing adeno-associated virus (AAV) core and a degradable polymeric shell that encapsulates MCL-1 siRNA (BIM/MCL-1 ChNPs) synergistically and selectively killed BCR-ABL+ CML cells in combination with dasatinib. In a mouse CML model, the BIM/MCL-1 ChNPs and dasatinib combination therapy suppressed proliferation of BCR-ABL+ hematopoietic cells and prevented leukemic infiltration of organs. The synergistic anti-leukemic effect was further pronounced in an acute phase model of the disease. This study investigated a strategy of developing a versatile and tunable multimodal therapy assisted by a computational toolset that analyzes the molecular foundation of a disease and predicts therapeutic response. The interdisciplinary approach developed and validated in this study can be used in discovering new therapies for cancer and other diseases.

Graphical abstract
summarizing the study design and approach. A schematic representation of the integrated in silico and in vivo pipeline utilized in the study. The workflow begins with in silico simulations, including Boolean network modeling and protein-protein interaction (PPI) network analyses, leading to target discovery, optimized nanoparticle design, and validation in pathological contexts. This was followed by therapeutic efficacy assessments of BIM/MCL-1 ChNPs and their combination with dasatinib in BCR-ABL+ leukemia models in vitro and in vivo.

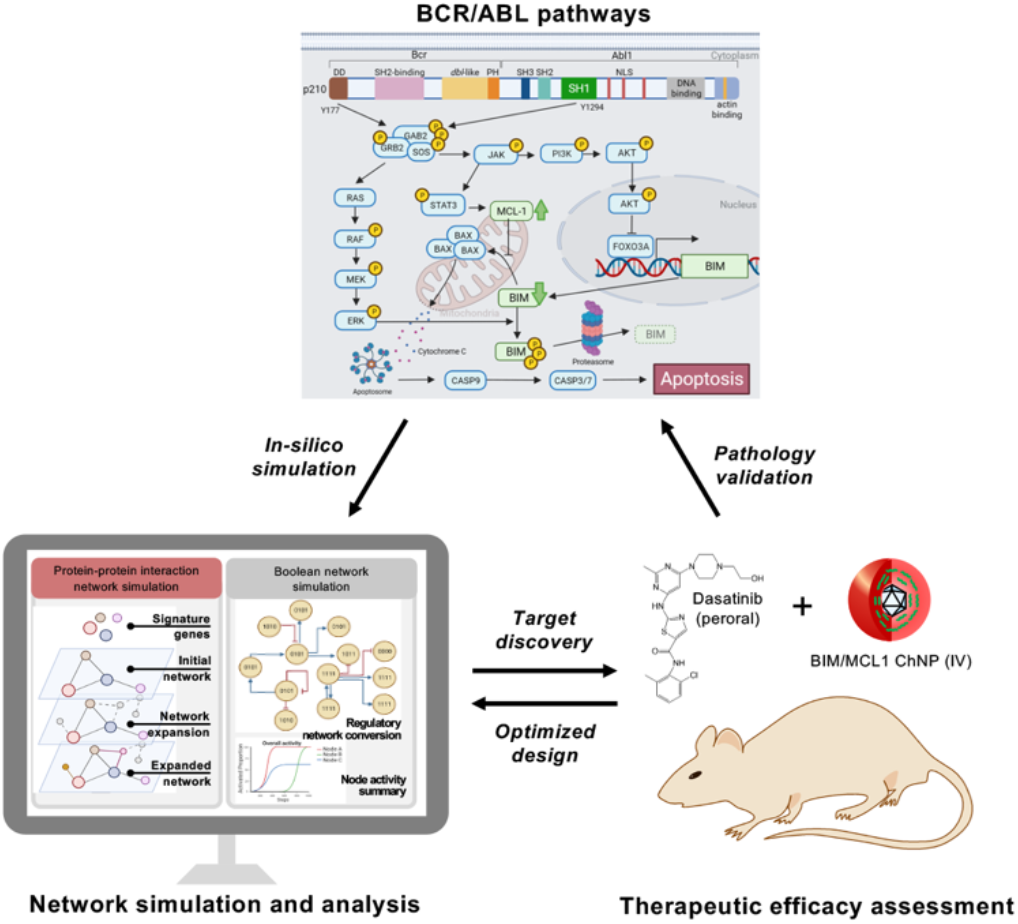

## Introduction

Cancer pathogenesis typically involves multiple aberrant, dynamically evolving cell signaling pathways (1). The pathological complexity and diversity of cancers, exacerbated by the myriads of drug resistance mechanisms, limit the efficacy of single-modal cancer therapies (2). This is evident in chronic myeloid leukemia (CML), where the clinical standard of care is tyrosine kinase inhibitors (TKIs). The development of the first BCR-ABL-targeted TKI, imatinib, was a breakthrough in CML treatment as it directly interfered with the enhanced kinase activity responsible for the activation of several pathways that promote cancer cell proliferation and impede apoptosis of BCR-ABL oncogenic fusion protein formed by a reciprocal translocation between chromosomes 9 and 22 (3–5). While imatinib was transformative in its approach to treating CML, its susceptibility to drug resistance necessitated the development of second-generation TKIs, including dasatinib, nilotinib, and bosutinib, as well as third-generation TKIs, including ponatinib, asciminib, and olverembatinib (6–7). Currently, the use of first and second generation TKIs, as frontline therapy, and third generation TKIs, as later-in line therapy, allows many individuals with CML to achieve long-term remission and near-normal life expectancy (8–10). However, a subset of patients is unable to tolerate TKI therapy or acquire drug-resistant mutations and experience treatment failure (11). This leads to acute phase progression and blast crisis, ultimately leading to mortality (12–13). Furthermore, insufficient eradication of leukemic cells is reported in approximately 50% of TKI-treated patients, thus the majority of CML patients must adhere to indefinite treatment (13–16). The adverse events associated with TKI use, though considered low-grade, are not inconsequential and include concerns for long-term organ toxicity (14–17).

The shortcomings in CML treatment highlight the need for a comprehensive, multi-dimensional therapeutic strategy that addresses the multiple molecular drivers enabling persistence in CML. While multi-drug regimens are typically sought out to compensate for the limitations of an individual drug, this approach is challenged by the large number of combination candidates, as well as the difficulty in conserving specificity and reducing the compounded off-target effects of each drug used (18–19). In this regard, computational analyses serve to comprehensively simulate the complex molecular mechanisms implicated in cancer-specific dynamics and can lead to the discovery of targets for combination therapy and predict therapeutic outcomes (20). The relatively well-documented pathways evaluated in immortalized cell lines and animal studies make CML an ideal model to demonstrate the capabilities of computational therapeutic assessment (21). Previously, synergized CML suppression was demonstrated through modulating BCR-ABL-dysregulated proteins by expressing the pro-apoptotic protein BIM and silencing the pro-survival protein MCL-1 (22). Notably, these key mediators in apoptotic pathways play critical roles in the efficacy of TKI therapy (23–25). In CML, the BCR-ABL oncogene suppresses the pro-apoptotic BIM and the restoration of its expression through TKI therapy has been recognized as a crucial mechanism of action with BIM levels serving as a prognostic biomarker in patients (26–29). Although BIM restoration alone is insufficient to induce the apoptosis of CML cells broad-spectrum inhibitors that silence anti-apoptotic proteins, including MCL-1, have been shown to sensitize CML cells to TKI therapy (30–32).

This study evaluates the hypothesis of inducing selective and synergistic apoptosis of CML by BIM-expressing, MCL-1 silencing viral/nonviral chimeric nanoparticles (BIM/MCL-1 ChNPs) in combination with a TKI, dasatinib, which is computationally simulated for target validation and efficacy predicted using *in vitro* and *in vivo* models. Computational construction and analysis of the protein-protein interaction (PPI) in CML employed a combination of network algorithms and Boolean networks to predict and assess the therapeutic outcomes of the combined genetic manipulation of BIM and MCL-1 by ChNPs and BCR-ABL by dasatinib. The therapeutic efficacy of co-administration of BIM/MCL-1 ChNPs and dasatinib was experimentally examined in CML murine cell lines and primary cells derived from patients at chronic phase and blast crisis of BCR-ABL+ (Ph+) CML and BCR-ABL-(Ph-) leukemia. The survival of mouse models recapitulating chronic phases and acute phases of CML were also assessed along with histological evaluations. This study not only provides an interdisciplinary, multimodal combined CML therapy development and validation pipeline, but also demonstrates new insight in developing efficient and safe therapies for other cancers and other diseases.

## Results

### Network-based simulation for understanding CML-specific signaling dynamics and therapy design with BIM/MCL-1 modulation and tyrosine kinase inhibition

To advance the understanding of CML-specific signaling dynamics, this study employed systems biology techniques, leveraging protein-protein interaction (PPI) networks and network algorithms to model the complex molecular interactions in CML and simulate BIM expression restoration and MCL-1 silencing. PPI networks connect proteins by their physical and functional interactions with edges and coupled with various algorithms to identify crucial proteins by their topology (e.g., centrality algorithms) or disease-association (e.g., Random Walk Restart [RWR]) (33–39). These approaches have demonstrated efficacy and utility in studying disease biology, including the discovery of novel targets and signaling pathways in gastric and esophageal cancers, as well as drug repurposing effectors for SARS-CoV-2 (40–42). Here, wild-type (WT) and CML networks based on a CML master gene list and the interactions of BCR-ABL (Fig. 1*A*) were constructed, followed by employing RWR to identify BIM and MCL-1 interactomes, thus elucidating their roles in apoptotic activity in CML through pathway enrichment (Fig. 1*A* and *SI Appendix* S1). The effect of BIM deletion polymorphism in CML was then simulated by computationally removing the BIM interactome (Fig 1*A*). Despite the simulation led to the removal of some apoptotic-related interactions in BIM and MCL-1 interactome (*SI Appendix* S1), the resulting networks demonstrated significant effects on apoptosis-related pathways in CML, highlighting reduced apoptotic activation in untreated CML compared to WT (Fig. 2*A*). To assess the functional consequences of pharmacological interventions, dasatinib interactors were curated from STITCH and DrugBank and modeled by eliminating dasatinib-interacting proteins from the networks (Fig. 1*B* and *SI Appendix* S2) (43–44). Notably, the combined modulation of BIM enrichment and MCL-1 silencing with dasatinib synergistically enhanced apoptotic pathway activation (Fig. 2*A* and *B* and *SI Appendix* S3), supporting their key regulatory roles in BCR-ABL-driven leukemogenesis.

**Figure 1.**
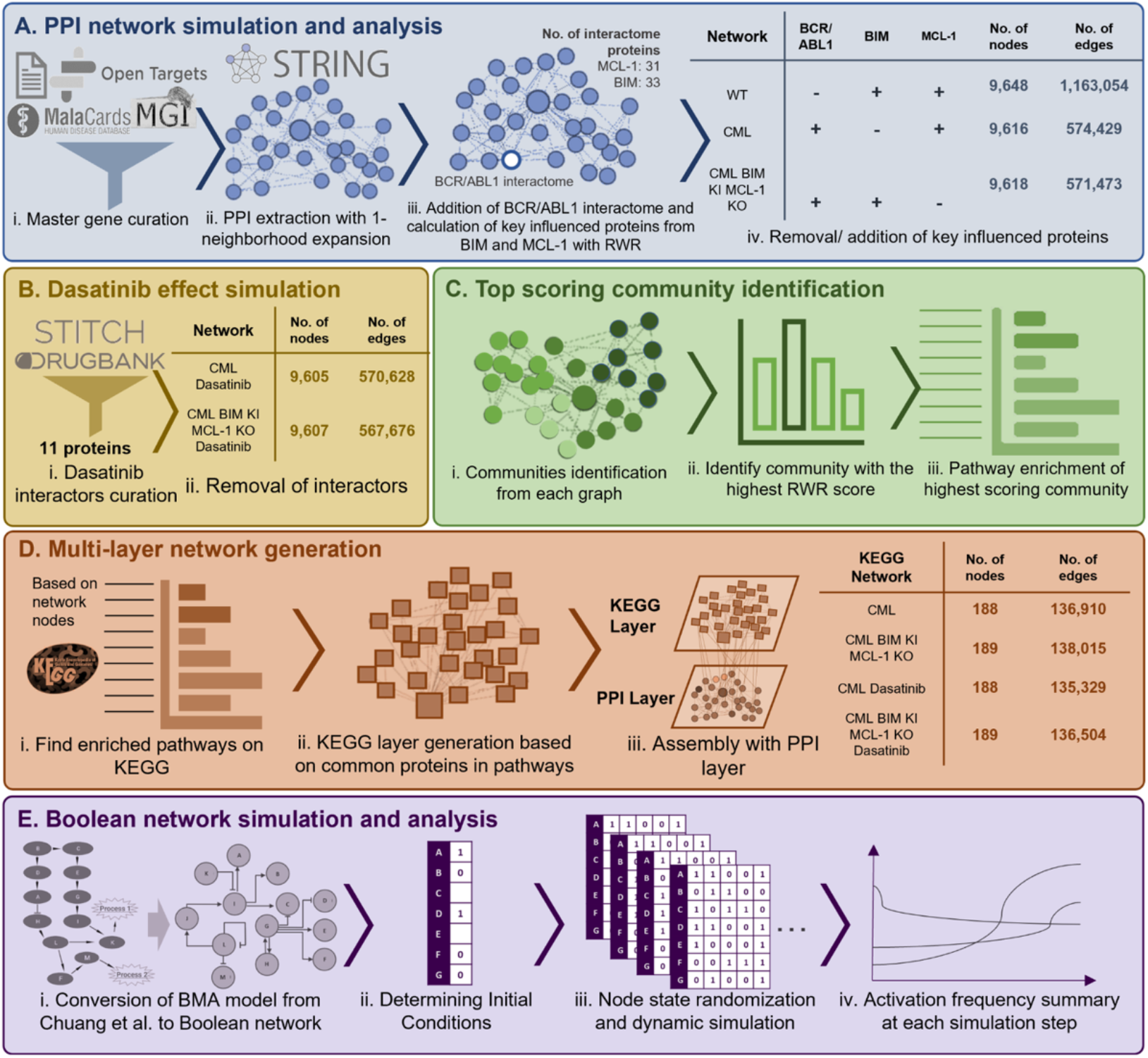
Construction and analysis of network models for dasatinib and BIM/MCL-1 ChNP effects in CML. (A) Network generation: (i) Master gene list curation from Open Targets, STRING, MalaCards, and MGI databases. (ii) PPI network construction and neighborhood expansion of BCR-ABL interactome. (iii) Addition of interactome proteins influenced by BIM and MCL-1. (iv) Generation of networks representing WT, CML, and CML conditions with BIM KI and MCL-1 KO, with node/edge counts indicated. (B) Dasatinib effect simulation: (i) Dasatinib interactors curated from STITCH and DrugBank. (ii) Effects of interactors removed in CML networks with and without BIM/MCL-1 modifications. (C) Top-scoring community identification: (i) Identification of communities from each graph. (ii) Selection of communities with the highest PageRank scores. (iii) Pathway enrichment analysis of the top-scoring communities. (D) Multi-layer network generation: (i) Enrichment analysis of KEGG pathways based on network nodes. (ii) KEGG layer generation using common pathway proteins. (iii) Integration of KEGG and PPI layers to form a comprehensive multi-layer network, with node/edge counts detailed for each condition. (E) CML Boolean network simulation: (i) Conversion of Bio Model Analyzer (BMA) network from Chuang et al. to Boolean network. (ii) Determination of initial conditions based on disease state and treatment strategies. (iii) Dynamic simulation with node state randomized at 2,000 steps and 5,000 randomized conditions. (iv) Summary of activation frequencies based on percentage of activated nodes at 5,000 randomized conditions.

**Figure 2.**
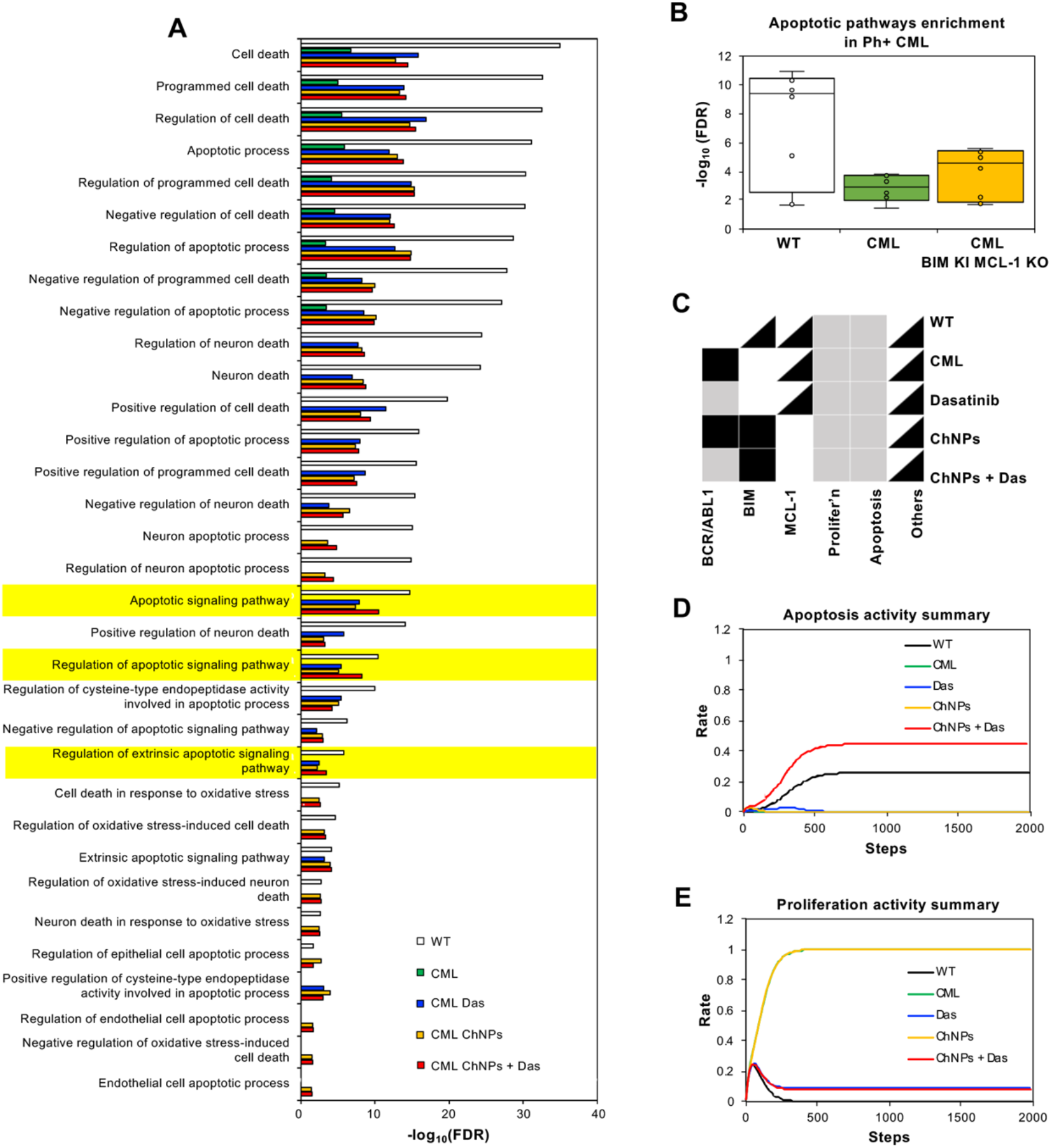
Computational simulation of BIM/MCL-1 ChNPs in alone and in combination with dasatinib with PPI and Boolean network. (A) PPI simulation showed that dasatinib and BIM/MCL-1 ChNPs, alone and in combination, restored the decreased association of BCR-ABL to apoptosis-related pathways in CML compared to WT network. (B) Summary of apoptotic pathways enrichment from PPI simulation indicated the restoration of apoptotic activities in BIM/MCL-1 ChNPs network from CML. (C) The node activation patterns in the CML Boolean network. Only the nodes that possess different activation patterns across simulation states are shown. White, grey, black and black-and-white boxes represent inactivation, partial activation, full activation, and randomized state, respectively. The activation frequencies of (D) Apoptosis and (E) Proliferation were observed across different simulation states.

Further refinement of the network simulation using community detection algorithms (Fig. 1*D* and *SI Appendix*, Fig. S4A) identified the level of apoptotic enrichment in top BCR-ABL scoring functional communities. Louvain community detection coupled with RWR scoring prioritized protein clusters with strong connectivity to BCR-ABL and/or BCR and ABL (*SI Appendix*, Fig. S4A). These analyses highlighted a significant shift in apoptotic signaling, with a more robust activation profile observed in conditions where BIM expression was restored and MCL-1 was silenced in combination with dasatinib. An additional analysis simulating BIM KO and MCL-1 KO also highlights the importance of BIM KI in the effect on enhanced apoptotic pathway activities (*SI Appendix*, Fig. S4B and S5). A multi-layer network incorporating KEGG pathway enrichment was generated to evaluate broader functional implications (Fig. 1*D*) (45–46). Enriched KEGG pathways were integrated with the PPI network to create a multi-layer heterogenous network model, where shared proteins were mapped across in connecting the enriched pathways. The multi-layer network results demonstrated that untreated CML had significantly lower apoptosis-related pathway activity, by RWRH scores, compared to WT, whilst introducing BIM/MCL-1 modulation with dasatinib significantly enhanced apoptotic activation (*SI Appendix*, Fig. S6). The comprehensive pathway-level assessments from the multi-layer network simulations align with the observations from both RWR-based and community-based PPI simulation results, revealing apoptotic and proliferative pathway shifts in CML under various treatment conditions.

### Boolean Network simulation predicts synergized apoptotic effect of BIM/MCL-1 ChNPs with dasatinib in CML

An extended evaluation was performed using Boolean network simulations to characterize the dynamic regulatory interactions within the CML network (Fig 1*E*) (47–48). The Boolean network model of CML which consists of 42 nodes (37 genes and 5 biological processes) and 65 regulatory interactions provided a framework to assess the frequency of apoptotic activation under distinct therapeutic conditions (*SI Appendix*, Fig. S7*A*). For each simulation, fixed states of BCR-ABL, BIM, and MCL-1 were assigned, based on the tested conditions (Fig. 2*C*), and their effects on each node were characterized by measuring its frequency being in the state of 1 (activation frequency) across 5,000 randomized networks. When BCR-ABL remained active in CML (green), apoptotic signaling was reduced, and proliferative pathways were upregulated, supporting the oncogenic role of BCR-ABL in CML progression (Fig. 2 *D* and *E*). However, transient BCR-ABL inactivation combined with sustained BIM activation and MCL-1 inhibition significantly enhanced apoptotic activation while suppressed proliferation, indicating a synergistic therapeutic potential for this multimodal approach (Fig. 2 *D* and *E*; red). Further functional importance validation by modeling the independent effects of individual BIM knock-in (BIM AAV), BIM knockout and MCL-1 knockout (MCL-1 siRNA) in CML resulted in distinct changes in apoptotic pathway regulation, highlighting the compensatory mechanisms that may arise upon individual modulation of these targets (*SI Appendix*, Fig. S5 and S7B-E). Overall, the computational results strongly support the therapeutic rationale for targeting BIM and MCL-1 alongside tyrosine kinase inhibition to achieve maximal apoptotic activation in CML.

### Synergized apoptosis of Ba/F3 p210 cells by BIM/MCL-1 ChNPs with dasatinib

ChNPs designed to simultaneously express BIM and silence MCL-1 were used for *in vitro* validation of the multimodal therapeutic approach. The ChNPs utilize the advantages of both viral and nonviral vectors by encapsulating a BIM-encoding AAV core in an acid-degradable polymeric shell that co-encapsulates siRNA targeting MCL-1 (BIM/MCL-1 ChNPs). The polymeric shell shields the viral core from the immune response and breaks down in the mildly acidic environment of the endosome to release siRNA and AAV (*SI Appendix*, Fig. S8). The genetic modulation by BIM/MCL-1 ChNPs with and without dasatinib co-administration was evaluated in the murine pro-B cell line Ba/F3 stably expressing BCR-ABL (p210 isoform). MCL-1 silencing was observed at 48 h after the treatment prior to BIM expression at 72 h (Fig. 3*Aii*). The time-dependent nature was likely a function of the ChNPs design, where acid-degradation of the polymeric shell in the endosome/lysosome resulted in the cytosolic release of siRNA, followed by AAV being trafficked to the nucleus for BIM expression. Though the dasatinib treatment alone did not exhibit an increase in BIM expression, the combined treatment of BIM/MCL-1 ChNPs and dasatinib upregulated BIM more than the BIM/MCL-1 ChNPs alone, supporting the dasatinib-promoted BIM restoration (49). MCL-1 levels were decreased by dasatinib to an even greater extent than the BIM/MCL-1 ChNPs, with similar levels observed using the combined treatment, likely approaching the limits of silencing by siRNA. Together, these results provided mechanistic evidence for synergized therapy by combining gene and targeted drug therapy for CML.

**Figure 3.**
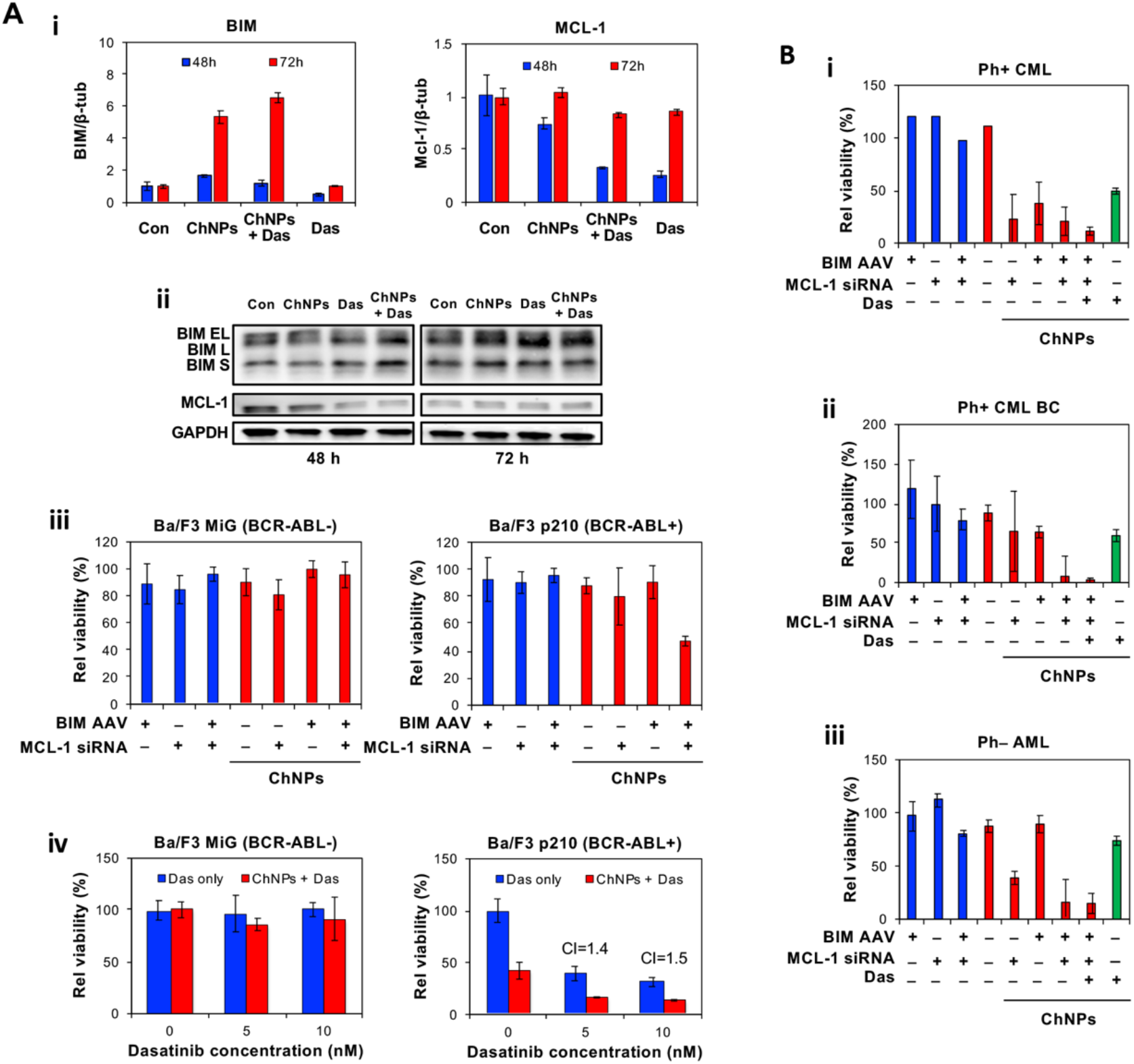
*In vitro* analysis of BIM/MCL-1 ChNPs alone and in combination with dasatinib on Ba/F3 wild type and BCR-ABL+ cell lines. (A) (i) BIM/MCL-1 ChNPs upregulated BIM RNA and downregulated MCL-1 RNA in Ba/F3 p210 (BCR-ABL+) cells, where combining BIM/MCL-1 ChNPs with dasatinib synergized the effect. (ii) BIM/MCL-1 ChNPs, dasatinib, and the combination of the two upregulated the protein expression of BIM and downregulated the protein expression of MCL-1 in Ba/F3 p210 (BCR-ABL+) cells. (iii) BIM/MCL-1 ChNPs did not affect BCR-ABLl-Ba/F3 cell lines, but specifically targeted BCR-ABL+ Ba/F3 cells for apoptosis (n=3). (iv) Combination of BIM/MCL-1 and dasatinib had no effect on BCRL-ABL-cells but act synergistically to killed BCR-ABLl+ leukemia cells (n=3). All error bars denote standard deviation. For (iii) and (iv) tables S1-3 detail *p*-values for all groups. (B) Effect of BIM/MCL-1 ChNPs on patient-derived primary leukemia cell viability. (i) ChNPs that contained BIM AAV, MCL-1 RNA, or the combination significantly decreased the viability of Ph+ CML patient derived cells, and the combination of dasatinib and BIM/MCL-1 ChNPs was more effective than any other treatment (n=3). (ii) BIM/MCL-1 ChNPs alone and combined with dasatinib had a significant effect on the viability of Ph+ CML patient-derived primary cells during blast crisis (BC) (n=3). (iii) ChNPs containing MCL-1 siRNA significantly decreased the viability of Ph-AML patient-derived primary cells, with a greater effect being observed with the combination of BIM AAV and MCL-1 ChNPs, dasatinib had no effect on the Ph-cells (n=3). Error bars denote standard deviation. All significance data detailed in tables S4-S6.

BIM and MCL-1 modulation by the ChNPs only affected cancer cells, as demonstrated by a viability study using Ba/F3 p210 and wildtype Ba/F3 empty vector (MiG) cells. The significant reduction in viability of the Ba/F3 p210 cells in contrast to the relatively unaffected viability of the Ba/F3 MiG cells by BIM/MCL-1 ChNPs demonstrated that this combination of targets impacts the dysregulated apoptotic signaling exhibited only by cancerous cells (Fig. 3*Aiii*). BIM/MCL-1 ChNPs outperformed all other treatment groups, including those incubated with free AAV and siRNA, indicating the importance of co-delivery for synergized apoptosis. The addition of dasatinib with the BIM/MCL-1 ChNPs resulted in a dasatinib dose-dependent decrease in the cell viability of Ba/F3 p210 cells, while Ba/F3 MiG cells were unaffected (Fig. 3*Aiv*). This result confirmed the improved effects of combined therapy *via* reprogramming cell signaling pathways to promote apoptosis as predicted by computational analyses with assured pathology-dictated specificity.

### Synergistic effect of BIM/MCL-1 ChNPs and dasatinib on primary patient leukemia cells

Primary cells isolated from patients diagnosed with BCR-ABL+ leukemia were treated with BIM/MCL-1 ChNPs in combination with dasatinib to assess the clinical potential of this therapy. The leukemias tested included BCR-ABL fusion protein (Ph)-positive chronic phase CML which is typically well-managed by TKIs and patients with blast crisis (BC) CML which is characterized by additional genetic alterations and a far worse prognosis (50–51). Additionally, this therapy was tested on cells isolated from acute myeloid leukemia (AML) that lacked the BCR-ABL translocation (Ph-) and are therefore not treatable by dasatinib. For each of the Ph+ CML cells, IC_50_ values of dasatinib were determined through dose escalation studies and used to test the combination of dasatinib with BIM/MCL-1 ChNPs compared to dasatinib alone (data not shown). Chronic Ph+ CML cells treated with ChNPs that contained BIM AAV alone, MCL-1 siRNA alone, or the combination of the two showed similar levels of cell viability, indicating that the ChNPs with either target was sufficient to achieve eradication of patient-derived Ph+ CML cells (Fig. 3*Bi*). In contrast, Ph+ BC CML cells required the combined targeting of BIM and MCL-1 to achieve eradication, demonstrating the benefit of regulating multiple effectors of the apoptotic pathways, especially during an elevated proliferation stage (Fig. 3*Bii*). In both the Ph+ CML cells, one time injection of ChNPs outperformed daily dasatinib treatment, achieving a relative viability level of ~20% in the chronic CML cells and ~8% in the BC CML cells, compared to ~40% and 60%, respectively. Using ChNPs and dasatinib in combination reduced cell viability further to ~10% in the chronic CML cells and ~3% in the BC CML cells. The combination therapy showed potential to fully eradicate leukemic cells, potentially preventing their recurrence. ChNPs were able to sensitize BC CML to dasatinib therapy, offering a novel therapeutic option for patients with traditionally hard-to-treat leukemias. The viability of Ph-AML cells was significantly reduced by BIM/MCL-1 ChNPs but only marginally affected by dasatinib due to their lack of the BCR-ABL oncogene (Fig. 3*Biii*). This effect is likely explained by the fact that many oncogenic pathways result in a downregulation of BIM and an upregulation of MCL-1 (27). The apoptotic efficacy of the BIM/MCL-1 ChNPs against the Ph-AML patient cells presents the possibility of developing synergistic therapies for Ph-negative by BIM-MCL-1 ChNPs in combination with a corresponding targeted drug, such as JAK or MEK inhibitors.

### BIM/MCL-1 ChNPs in combination with dasatinib extended survival and delayed disease progression in chronic and acute phase CML leukemia in mice

A transduction-transplantation mouse model of CML was used to test the efficacy of BIM/MCL-1 ChNPs in combination with dasatinib *in vivo* (Fig. 4A) (52). The p210+ cells co-expressed GFP to enable monitoring of the CML burden by tracking the percentage of GFP+ leukocytes. For the studies of chronic phase CML, BCR-ABL+ cells in the blood at the initial point post-transplant time point ranging from 30-40% whereas acute phase CML was modeled by higher starting levels of 50-60% BCR-ABL+ cells. In both disease models, BIM/MCL-1 ChNPs were administered via intravenous tail injection once on day 0 for the chronic model and twice on days 0 and 7 for the acute model while dasatinib was administered by daily oral gavage until withdrawn at day 40 (Fig. 4*A*). In addition to assessing survival, the mice were monitored for disease burden (GFP+ cells in peripheral blood leukocytes), white blood cell count, and body weight for the duration of the study (*SI Appendix*, Fig. S9 and S10). In the chronic model, a one-time treatment with BIM/MCL-1 ChNPs alone was not sufficient to fully halt disease progression. However, even a single dose of BIM/MCL-1 ChNPs extended survival of CML mice. Mice in the untreated group succumbed to the disease by day 97, while the ChNPs-treated group survived until day 163 (Fig. 4*B*). Prior to withdrawal of the dasatinib administration, 100% of the mice treated with the combination treatment of BIM/MCL-1 ChNPs and dasatinib survived, compared to 78% with dasatinib alone. Additionally, survival until the end of the study was observed in 29% of the mice receiving the combined treatment, compared to 14% of those treated with dasatinib alone. These results suggested that the BIM/MCL-1 ChNPs had a vital role in slowing disease progression. The mice in the acute CML model with higher leukemic burden succumbed to disease faster than in the chronic model (Fig. 4*C*). To respond to this severity, a second administration of BIM/MCL-1 ChNPs was given one week following the initial dose, while daily oral administration of dasatinib was continued. In this model, the BIM/MCL-1 ChNPs were found to extend the survival of treated mice, compared to the untreated group. Only 25% of mice survived until the withdrawal of dasatinib in the group treated with dasatinib alone, compared with 75% of mice in the dasatinib and BIM/MCL-1 ChNPs combination group. However, due to the severity of the disease, mice in both the dasatinib-only and the combination treatment groups quickly succumbed to the disease after dasatinib withdrawal. These results support the role of BIM/MCL-1 ChNPs in sensitizing advanced and aggressive CML to TKIs, playing even more critical roles in suppressing the acute state than the chronic state of the disease.

**Figure 4.**
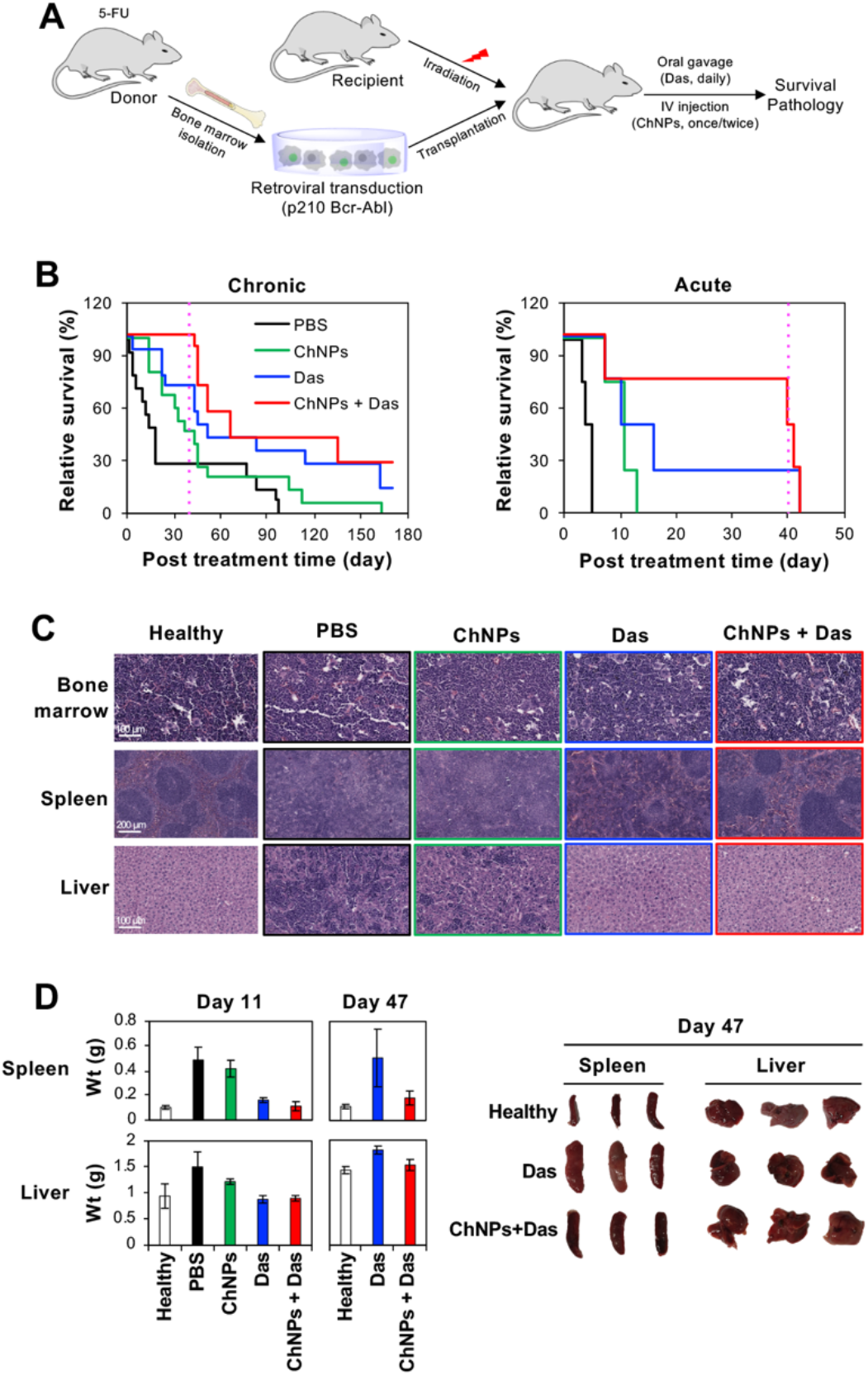
In vivo BCR-ABL+ leukemia model treated with BIM/MCL-1 ChNPs and dasatinib. (A) Transduction-transplantation model to induce naturally progressing BCR-ABL+ leukemia in C57BL/6 mice. (B) In both chronic (n=12 for PBS and ChNPs groups, n=14 for dasatinib and combination of dasatinib and BIM/MCL-1 ChNPs group) and acute (n=4 per group) models of BCR-ABL+ leukemia, BIM/MCL-1 ChNPs alone extended average survival compared to PBS, and the combination of BIM/MCL-1 ChNPs and dasatinib extended life before dasatinib treatment was stopped. In the chronic model, after dasatinib treatment was ended, mice had a longer average survival, with 29% of mice surviving until the end of the experiment compared to 14% of mice treated only with dasatinib. (C) Histology on mouse bone marrow, spleen, and liver displayed that dasatinib and BIM/MCL-1 ChNPs combined allowed for the structure to be similar to that in healthy mice. PBS, BIM/MCL-1 ChNPs, and dasatinib treated mice had high infiltration of white blood cells and structural organ damage. (D) Spleen and liver weights and images of mice with various treatments (n=3 per group). Dasatinib alone and in combination with BIM/MCL-1 ChNPs were effective at keeping liver and spleen weights aligned with healthy mice, but after dasatinib withdrawal, only BIM/MCL-1 ChNPs in combination with dasatinib treated mice were able to maintain healthy sized livers and spleens. All error bars are standard deviation. See tables S7-S12 for significance data on all experiments in this figure.

The impact of BIM/MCL-1 ChNPs with and without dasatinib on the bone marrow, spleen, and liver was evaluated through histopathology and weight changes (Fig. 4*C* and *D*). Histological analysis revealed a predominance of neutrophils in the bone marrow, obliteration of splenic architecture, and infiltration of myeloid cells into the liver in all mice except those treated with ChNPs and dasatinib in combination. Combination therapy protected the histological architecture of the organs in the CML mouse model to a degree comparable to that of the healthy mice, indicating reduced disease manifestation. Additionally, the liver and spleen weights demonstrated that both dasatinib and the combined ChNPs and dasatinib treatments achieved a reduction in myeloid infiltration at day 11. In contrast, the combined treatment exhibited an even greater reduction than treatment with dasatinib alone at day 47. Taken together, these results demonstrate the potential clinical advantage of combined CML treatment with BIM/MCL-1 ChNPs and dasatinib to suppress leukemic proliferation.

## Discussion

The rational development of efficient cancer therapies *via* a mechanistic understanding of cancer signaling pathways has been limited due to unexpected crosstalk, redundancy, and activation of compensatory signaling (18). Additionally, the vast combinations of pathway combinations and specific cancer type applications exceed the capacity for study, resulting in dispersed efforts and stagnant clinical translation (17–18). The computationally guided approach of molecularly reinforcing targeted drug activity with dual targeting of apoptosis genetic mediators directly confronts these challenges. This study predicted the facilitated apoptosis of CML cells by simultaneous expression of pro-apoptotic BIM and silencing of anti-apoptotic MCL-1 could be further synergized by the BCR-ABL TKI, dasatinib (Fig. 3*Aiv*). The synergistic effect which was predicted through computation and achieved when combining the use of dasatinib further supported that this dual modulation was able to overcome the dysregulated apoptotic state responsible for limiting the molecular response achieved by the targeted drug (Fig. 2*D* and *E*). The significance of this capability is especially evidenced by the pronounced improvement observed in the combination treatment in a CML model that is not well managed by the use of dasatinib alone which included patient-derived BC CML cells and an *in vivo* model with a high burden of leukemic cells representing the acute phase of the disease (Fig. 4*B*). The outcomes here lend support to the goal of enabling targeted drugs to achieve effective eradication that would eliminate persistence, thereby removing the harms of prolonged drug use and risks of acquiring untreatable drug resistance. The approach developed and validated in this study is readily applicable to developing therapies for other cancers and other diseases treated with targeted therapy. It is also expected to continually improve in accuracy and reliability alongside the rapid advances in computational toolsets, accumulations of multi-molecular, spatial, and single cellular databases, and emerging therapeutic modalities.

Cancer-targeted therapy has been attempted by accumulation of therapeutic agents in cancerous tissue, specific binding to cancer cells, and selective activation by cancer-specific stimuli, yet has had minimal success using these avenues (53). A remarkable advantage of the dual targeting of oncogenic pathways that facilitate cellular apoptosis is its specific biological impact on cancer cells, which bypasses the need for the aforementioned strategies. The observation that simultaneous BIM overexpression and MCL-1 silencing by the ChNPs did not affect the viability of non-cancerous cells but effectively killed CML cells demonstrated a distinct vulnerability caused by the imbalance of the apoptotic mediators induced by upstream oncogenic drivers (Fig. 3*Aiii*). Cancer therapeutics acting in a pathologically determined manner rather than relying on targeted delivery and activation is technologically feasible with high specificity. Regardless of the particular upstream driver of the aberrant signaling that causes the dysregulation, it is the resulting imbalance of pro-apoptotic and anti-apoptotic mediators that ultimately enables survival and evasion of cancer cell death (54–55). As such, the overexpression of BIM and silencing of MCL-1 serve to subvert oncogenic signaling to molecularly ensure the promotion of apoptosis directly at the point of execution (54–56). Though the levels of BIM and MCL-1 in particular have been implicated in the efficacy of targeted cancer drugs, this combined approach has not been feasible using small molecule drugs due to challenging normalization of their pharmacokinetics characterization (57–59). Adding a targeted drug, such as dasatinib against BCR-ABL, further augments the pathologically specific therapeutic outcomes. Manipulating the primary molecular driver in concert with the underlying effector pathways not only synergizes therapeutic efficacy but also minimizes the chance of developing resistance by circumventing oncogenic signaling through the uses of multiple apoptotic enforcements. To strengthen these findings, further studies could be conducted in examining the dose and treatment scheduling regimen of ChNPs required to extend survival while reducing the need for the targeted drug altogether. The use of multiple administrations of BIM/MCL-1 ChNPs can be evaluated for its impact on residual cancer cell populations not fully eradicated by prolonged targeted-cancer drug treatment alone.

This BIM/MCL-1 ChNPs and dasatinib combinatorial treatment study demonstrates the potential for a widely applicable approach to improving targeted therapy. While the focus of this study was CML, these results also provided valuable indications for further use of our combination targeted strategy for other cancers and diseases. In the evaluation of patient-derived AML cells which lack the BCR-ABL driver and was therefore anticipated to not be responsive to dasatinib treatment, it was notable that BIM and MCL-1 modulation alone resulted in cancer cell death. This exemplified how direct targeting of apoptosis in downstream pathways can be effective even for cancers with signaling pathways that are not well understood and lack a druggable molecular driver. In addition to its use in highly aggressive and heterogenous leukemias, the *in vivo* delivery capability of the ChNPs in reaching leukemic cells established in the bone marrow warrants evaluation for its use in solid tumors. The extended applications of the ChNPs are also further enabled by the capabilities of the computational approach used in this study. In developing the model of CML signaling interactions in the context of the proposed combination targeting therapy, a versatile methodology as established for pathway construction and analysis that can be applied to various cancers (Fig. 1). This enables the evaluation of various phenotypes in their full molecular context for comprehensive mechanistic insights of the key interactions that contribute to pathogenesis and drug efficacy. As a tool for *in silico* therapeutic screening, these models can accelerate the process of optimal combinatorial target discovery and clinical translation. Taken together, the interdisciplinary strategy demonstrated in this study creates a universally tunable cancer therapy that addresses the current unmet clinical challenges in developing efficient and safe combination therapies for cancer.

## Materials and Methods

### Construction of PPI Networks

A list of master genes was curated from various sources (detailed in Supplementary Information) and used to extract the protein-protein interactions (PPIs) from the STRING database (version 11.0) with expansion by one neighborhood (directly interacting proteins) (36). The BCR-ABL interactome was added from the data published in Brehme et al. (21). For the CML network, the common deletion polymorphism of Bcl-2 like protein 11 (BIM) in CML was recreated by removing BIM and its associated nodes (Random Walk Restart result (RWR; p-value ≤ 0.01) starting from BIM; 33 nodes) from the resultant multi-omics network. For the WT network, BCR-ABL and the associated interactions were removed instead. To simulate the CML biology under BIM/MCL-1 ChNPs treatment, MCL-1 and its associated nodes (RWR result (p-value ≤ 0.01 at 100 permutation) starting from MCL-1; 31 nodes) were removed from the resultant PPIs. For dasatinib effect simulation, the dasatinib targets from STITCH and Drug Bank were removed from the simulated networks (43–44).

### Key nodes identification and functional enrichment analysis

#### PPI Network Permutation

In each network, edge betweenness centralities were calculated and used as edge weights before performing RWR using igraph (36). For each network, RWR starting from BCR, ABL, and BCR-ABL (if available) was performed to identify the key nodes. Degree, betweenness, and eigenvector centrality scores were also calculated using NetworkX37. The top influential bides were subsequently selected using the empirical p value (≤ 0.05 for centrality scores and ≤ 0.01 for RWR scores at 100 permutation) as a cut-off after 100 network permutations. To gain mechanical insights into the key nodes, g:Profiler was used to identify the enriched biological pathways with the key node lists (38). Fisher’s exact test was applied to compute the p-value of the level of enrichment for each pathway with Benjamini-Hochberg False Discovery Rate (FDR) correction.

Permutation tests were performed 100 times to identify significant nodes for the centralities and RWR algorithms applied on each multi-omics network. In each permutation test, a random network was generated with a preserved degree distribution of the original multi-omics network by reconnecting the edges and swapping the nodes. For each permutated network, the centrality and PageRank algorithm were applied to obtain the cumulative results used to calculate the empirical p value of the network algorithm. The four permutation test results were combined to determine the final set of key nodes that have an empirical p value of ≤0.05 in centrality scores and ≤ 0.01 for RWR scores in each network.

#### CML Boolean Network Simulation

CML regulatory elements were identified and detailed in Chuang et al. (47). The CML Qualitative Networks were simplied by “chain shrinking” (e.g. removing nodes that would have the same state as their regulators and pass the value to their affecter nodes) and no priorities were assigned between the regulations affecting the same node (e.g. all regulatory relationships were expressed as OR). The resultant Boolean network contains 42 nodes (5 biological processes: Apoptosis, Proliferation, Correct Differentiation, Growth Arrest, and Self-Renewal) and 65 regulatory edges. For each condition, node states dependent on the conditions were assigned (Fig. 2*C*) and 5,000 possible combinations were generated for the randomized states of other nodes. The model was updated with a general asynchronous updating method for 2,000 steps and the state of each node was recorded at each step.

### Preparation and characterization of AAV/siRNA ChNPs

ChNPs were formulated with polyketal monomers and cross-linkers that were synthesized as previously mentioned (22). Eosin-tagged AAV (eosin-AAV) was formed by incubating 6×10^10^ GC of null AAV2 or BIM (isoform L) AAV2 (Vector Biolabs, Malvern, PA) with eosin-5-isothiocyanate (Biotium, Fremont, CA) in 10 mM sodium bicarbonate overnight at 4ºC with gentle agitation. The next day, eosin-AAV2 was purified by filter centrifugation in 100kDa Amicon filters (Thermo Fisher Scientific, Waltham, MA), at 1600⋅g for 5 min at 4ºC. After Eosin-AAV was washed with 10 mM HEPES and filter centrifugation was repeated twice, 800 µL of 10 mM HEPES was added to a vial with a magnetic stir bar, followed by 2.5 mg monomer, 0.5 mg crosslinkers, 3 µg of scrambled siRNA ([Scr siRNA] ThermoFisher Scientific, Waltham, MA) or MCL-1 siRNA (Qiagen, Hilden, Germany), and 10 mg ascorbic acid in 200uL. Vials were placed on ice and stirred for 3 min. 6×10^10^ GC of eosin-AAV in 500 µL were added to the vial and stirred for 1 min. White light at 760 klux using a halogen lamp was applied for 10 min. Then 2.5 mg monomer and 1 mg crosslinker were added to the vial, and they were allowed to react for an additional 5 min in light. Light was turned off and vials were stirred for 1 min. in the dark before removing to isolate the BIM/MCL-1 ChNPs. ChNPs were collected and isolated by filter centrifugation using a 100kDa Amicon filter at 1,600×g for 30 min at 4ºC. ChNPs were washed with 10 mM HEPES and centrifugation was repeated twice, for a total of three times. Resulting ChNPs were incubated at 37ºC in acetic acid (pH=5.5) for 4 h to mimic endosomal conditions. Free AAV, intact NPs, and hydrolyzed ChNPs were then treated with 20 µg/mL RNAse A (Thermo Fisher Scientific, Waltham, MA), at 37ºC for 20 min, then 1 mg/mL proteinase K (Thermo Fisher Scientific, Waltham, MA), for 30 min at room temperature in order to lyse the AAV. AAV DNA was extracted using Wako DNA Extractor SP kit (Wako Chemicals, Richmond, VA) and AAV titer was determined using a QuickTiter™ AAV Quantitation Kit (Cell Biolabs, Inc, San Diego, CA). To determine siRNA encapsulation, protection, and release: free AAV, intact NPs, and hydrolyzed ChNPs were assessed using a RiboGreen™ RNA Assay Kit (ThermoFisher Scientific, Waltham, MA). Free siRNA, intact NPs, and hydrolyzed NPs were stained and visualized using agarose gel electrophoresis. ChNPs were collected and resuspended in 1 mL of 10 mM HEPES at a final concentration of 6×10^10^ GC/mL AAV and 0.75 µg/mL siRNA. ChNPs in 100 µL were combined with 600 µL of DI H2O for size and zeta potential analysis by dynamic light scattering (DLS) using a Zetasizer Nano (Malvern Instruments, Malvern, United Kingdom).

### Cytotoxicity by BIM/MCL-1 ChNPs and dasatinib

Ba/F3 MiG, a murine pro-B-cell line, and Ba/F3 p210 (Ba/F3 cells expressing p210 BCR-ABL fusion protein), primary patient CML, CML BC, and AML cells were obtained from Angela Fleischman (University of California, Irvine) and cultured in RPMI media (ThermoFisher Scientific, Waltham, MA), supplemented with 10% FBS (ThermoFisher Scientific, Waltham, MA) and 1% penicillin-streptomycin (ThermoFisher Scientific, Waltham, MA). BaF3 MiG cells were additionally supplemented with 10 ng/mL IL-3 (BioLegend, San Diego, CA) to support their proliferation. Primary patient cells were obtained from through IRB #2014-1709 (University of California, Irvine) and cultured in X-Vivo 20 media (ThermoFisher Scientific, Waltham, MA) supplemented with Flt-3, TPO, IL-3, and SCF (Biolegend, San Diego, CA). All cells were cultured at 37ºC, 5% CO2, and 100% humidity. In assessing the transduction and apoptosis, cells were seeded at 10,000 cells/well in a 96-well plate in 100 µL of their respective media. After a 24-h of incubation, 20 µL of 10 mM HEPES alone or containing 6×10^10^ GC/mL AAV and/or 3 µg/mL siRNA, both free, and encapsulated in ChNPs were added to the cells. After a further 24 h, media was replaced with 100 µL of fresh media. Then 48 h later, 10 µL of 3-(4,5-dimethyl-2-thiazolyl)-2,5-diphenyltetrazolium bromide (MTT [Sigma-Aldrich, Milwalkee, WI]) was added to each well. After 4 h of incubation, the media was aspirated and 100 µL of DMSO was added to cells. The plates were measured for absorbance at 570 nm using a Synergy HT (BioTEK, Winooski, VT, USA) microplate reader. To assess the cytotoxicity by dasatinib, cells seeded for 24 h at a density as described earlier incubated with a serially diluted dasatinib, with a maximum dose of 10 nM, with or without 1×10^10^ GC/mL BIM/MCL-1 ChNPs. Media was replaced after 24 h, and further incubated for 48 h before MTT assay was performed as described earlier.

### Western blot and real time polymerase chain reaction (RT-PCR)

Cells treated as described earlier were collected and lysed by incubating them in RIPA lysis buffer (ThermoFisher Scientific, Waltham, MA) for 30 min on ice. After cell debris pellet was removed by centrifugation for 20 min at 16,000×*g* at 4ºC, supernatant was harvested and run in a 12% SDS page gel before being transferred to a nitrocellulose membrane (Thermo Fisher Scientific). The membrane was blocked for 1 h at room temperature in a blocking buffer (Thermo Fisher Scientific). The primary antibodies against GAPDH (Cell Signaling Technology, Danvers, MA), MCL-1 (Cell Signaling Technology), and BIM (Thermo Fisher Scientific,) were diluted in 5% BSA at a 1:1,000 and incubated with the membrane overnight at 4ºC with slight agitation. The membranes were then rinsed three times with 1X TBST (Thermo Fisher Scientific) then incubated with secondary goat IgG anti-rabbit antibody (Thermo Fisher Scientific) diluted at 1:1,000 in TBST for 1 h at room temperature (Cell Signaling Technology), rinsed 3 times with TBST, and imaged using a GBox (Syngene, Frederick, MD). RNA from treated cells was extracted using Trizol reagent (Invitrogen, Carlsbad, CA) according to the manufacturer’s instructions. A total of 1 µg RNA was used for cDNA synthesis according to the manufacturer’s instructions (Bio-Rad, Hercules, CA). RT-PCR was carried out using a Quant Studio 7 (Thermo Fisher Scientific). Reactions were run in triplicate in three independent experiments. The geometric mean of housekeeping gene Beta tubulin was used as an internal control to normalize the variability in expression levels and was analyzed using the 2-ΔΔCT method.

### Anti-leukemic effects by ChNPs and dasatinib using a transduction/transplantation p210 mouse model

All animal experiments were approved by the Institutional Animal Care and Use Committee (IACUC) at the University of California, Irvine. C57BL/6 mice, 6-8 weeks old, originally obtained from Jackson Laboratory (Bar Harbor, ME) but bred in house. Donor mice were injected retro-orbitally with 150 mg/kg of 5-fluorouracil (5-FU) (American Pharmaceutical Partners, Schaumburg, IL) and allowed to rest for 5 days. Mice were then sacrificed, bone marrow was collected, and the bone marrow was transduced to express the p210 BCR-ABL transgene and GFP using 2×10^6^ PFU/mL of p210 expressing retrovirus (obtained from Professor Angela Fleischman). Recipient mice were irradiated at 800 rads to eradicate their bone marrow were injected with 5×10^5^ cells of 12% p210/GFP+ bone marrow from donor mice within 24 h, as well as 2×10^5^ whole bone marrow cells from a wild-type mouse for hematologic support during the recovery phase post-transplant. After 7 days transplanted mice were injected intravenously with 150 µL of PBS alone or 5×10^11^ GC BIM/MCL-1 ChNPs once on day 0 for chronic model and twice, once on day 0 then again on day 7 for the acute model. Half of each group was further treated daily with 20mg/kg dasatinib dissolved in 8 mM citrate buffer via oral gavage for 40 days in both disease models. Survival was tracked recording the date of death for each mouse over the 65 days following initial treatment. Mice were weighed weekly and recorded. Any mouse that lost more than 20% of its starting weight was euthanized. Blood samples were collected once a week and the GFP population was determined using flow cytometry (GUAVA EasyCyte) (Luminex Corporation, Austin, TX). Every two weeks mouse blood was further analyzed for complete blood counts using an ABCVet Hemalyzer (Scil, Altorf, France). After 10 days of treatment, half of each treatment group was sacrificed and the spleen, liver, and one femur were collected and examined for histology. In the double injection study, all of the above protocols were the same, with the addition of a second injection at the same dose to the primary treatment.

### Statistical analysis

All data are presented as mean ± standard deviation. Statistical analysis was performed using a two-tail Student’s t-test with a significance threshold of *p* < 0.05. For functional enrichment analysis, statistical analysis was performed using the Mann-Whitney U rank test on the −log10(FDR) between each network.

## Acknowledgments

We thank Melissa Thone for assisting with mouse experiments. This research is financially supported by the National Cancer Institute of the National Institute of Health (NIH/NCI) (1R21CA228099-01A1). The UCI Chao Family Comprehensive Cancer Center is supported by NIH/NCI (P30 5P30CA062203) and its Experimental Tissue Resource (ETR) shared resource is supported by the NIH/NCI (P30CA062203). R.T.L. was supported by the UCI Interdisciplinary Cancer Research T32 training grant awarded by the NCI (T32CA009054) and UCI Medical Scientist Training Program (MSTP). The content is solely the responsibility of the authors and does not necessarily represent the official views of the NIH. C.W.L. is supported by the Milner Therapeutics Institute and the Department of Surgery, University of Cambridge. N.H. is funded by LifeArc.

## Notes

### Competing Interest Statement

The authors have declared no competing interest.

