## Supplementary Information for "Multimodal gene and targeted drug therapy for chronic myelogenous leukemia: Computational target analysis and therapeutic validation"

[1]Department of Chemical and Biomolecular Engineering, University of California, Irvine, CA 92697, United States. [2]The Milner Therapeutics Institute, University of Cambridge, Cambridge, United Kingdom, CB2 0AW. [3]Department of Surgery, University of Cambridge, Cambridge, United Kingdom, CB2 0QQ. [4]Department of Biomedical Engineering, University of California, Irvine, CA 92697. [5]Department of Pharmaceutical Sciences, University of California, Irvine, CA 92697. [6]Department of Chemical Engineering and Biotechnology, University of Cambridge, Cambridge, United Kingdom, CB3 0AS. [7]Division of Hematology/Oncology, Department of Medicine, University of California, Irvine, CA 92697. 8Cambridge Centre for AI in Medicine, Department of Applied Mathematics and Theoretical Physics, University of Cambridge, Cambridge, United Kingdom, CB3 0WA. 9Cambridge Stem Cell Institute, University of Cambridge, Cambridge, United Kingdom, CB2 0AW. [10]Department of Molecular Biology and Biochemistry, University of California, Irvine, CA 92697.

[†] These authors equally contributed to the work.
[‡] Current address: Avatrial Ltd., Cambridge, United Kingdom, CB2 3QJ.
[¶] Current address: CardiaTec Bioscience Ltd., Cambridge, United Kingdom, CB2 1GE.

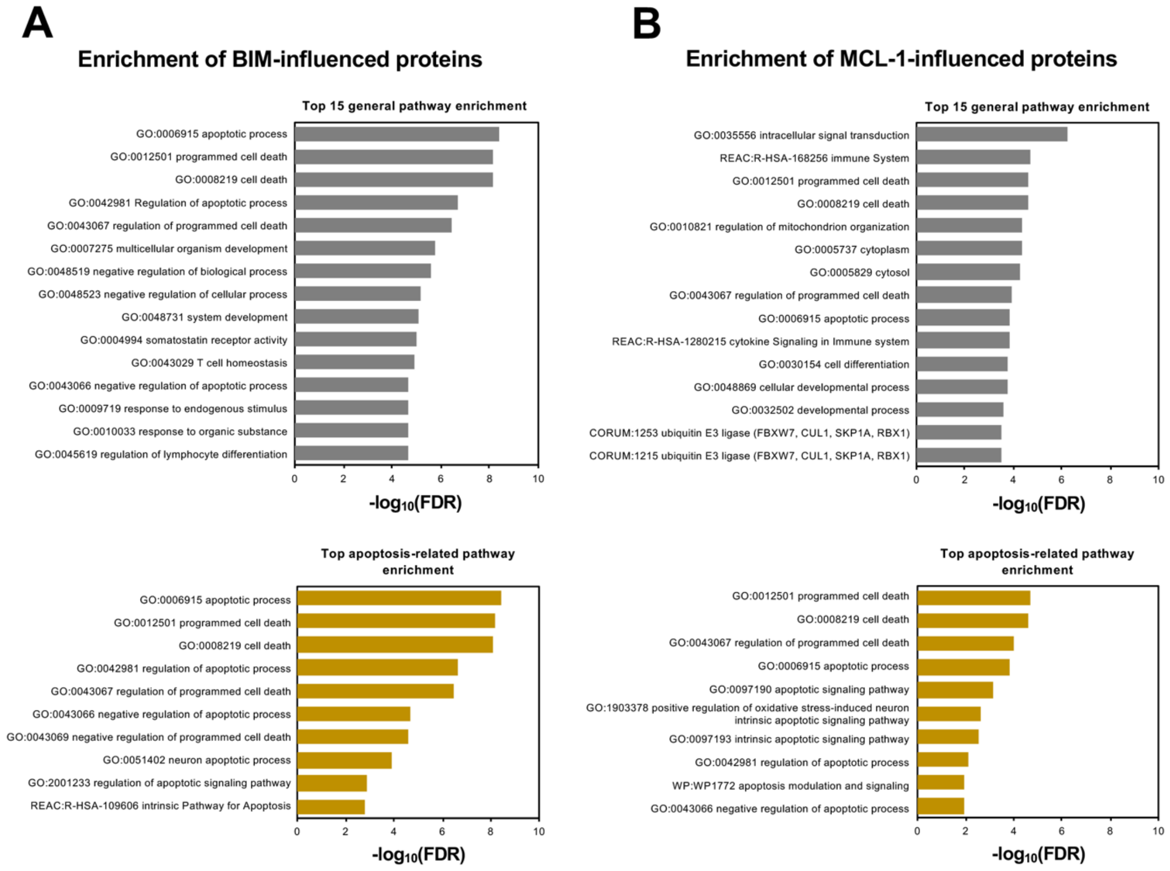


**Figure S1. Enrichment of BIM-influenced and Mcl1-influenced proteins.** Bar plots showing the -$\log_{10}$(False Discovery Rate) (FDR) of the top 15 pathways (top panel) and apoptosis-related pathways (bottom panel) enriched by the (A) BIM-influenced proteins identified from BIM Random Walk Restart (RWR) and (B) MCL-1-influened proteins from MCL-1 RWR in the CML PPI network. The high $\log_{10}$(FDR) of the apoptosis-related pathway suggests a large number of apoptotic-related proteins have been removed in CML BIM KI MCL-1 KO network, thus reducing the apoptosis pathway enrichment during BIM KI/MCL-1 KO ChNPs simulation.

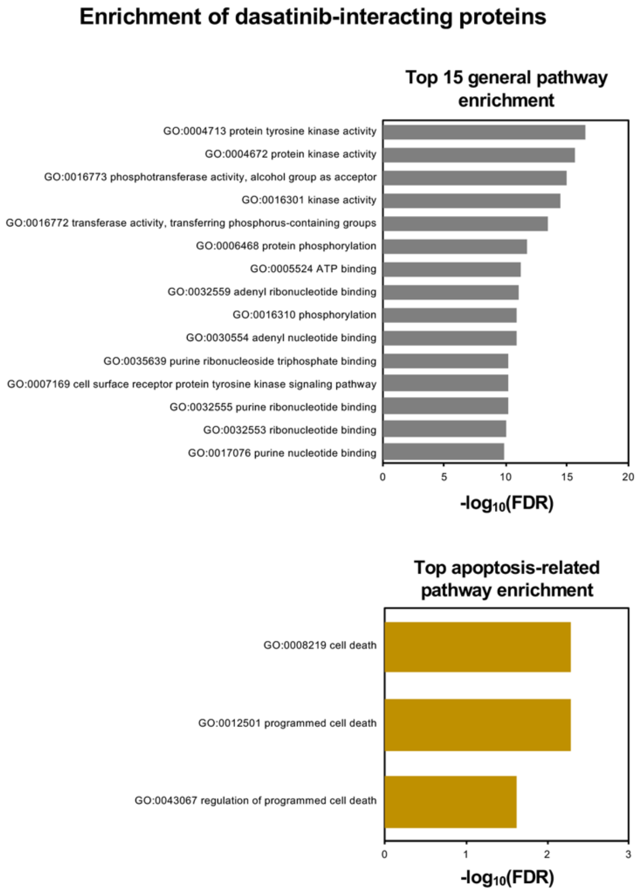


**Figure S2. Enrichment of Dasatinib interacting proteins.** Bar plots showing the $-\log_{10}$(False Discovery Rate) (FDR) of the top 15 general pathways (top panel) and apoptosis-related pathways (bottom panel) enriched by Dasatinib interacting protein. The high $\log_{10}$(FDR) of the apoptosis-related pathway suggests the removal of apoptotic-related proteins in CML BIM KI MCL-1 KO Dasatinib network, thus reducing the apoptosis pathway enrichment during BIM KI/MCL-1 KO ChNPs + Das simulation.

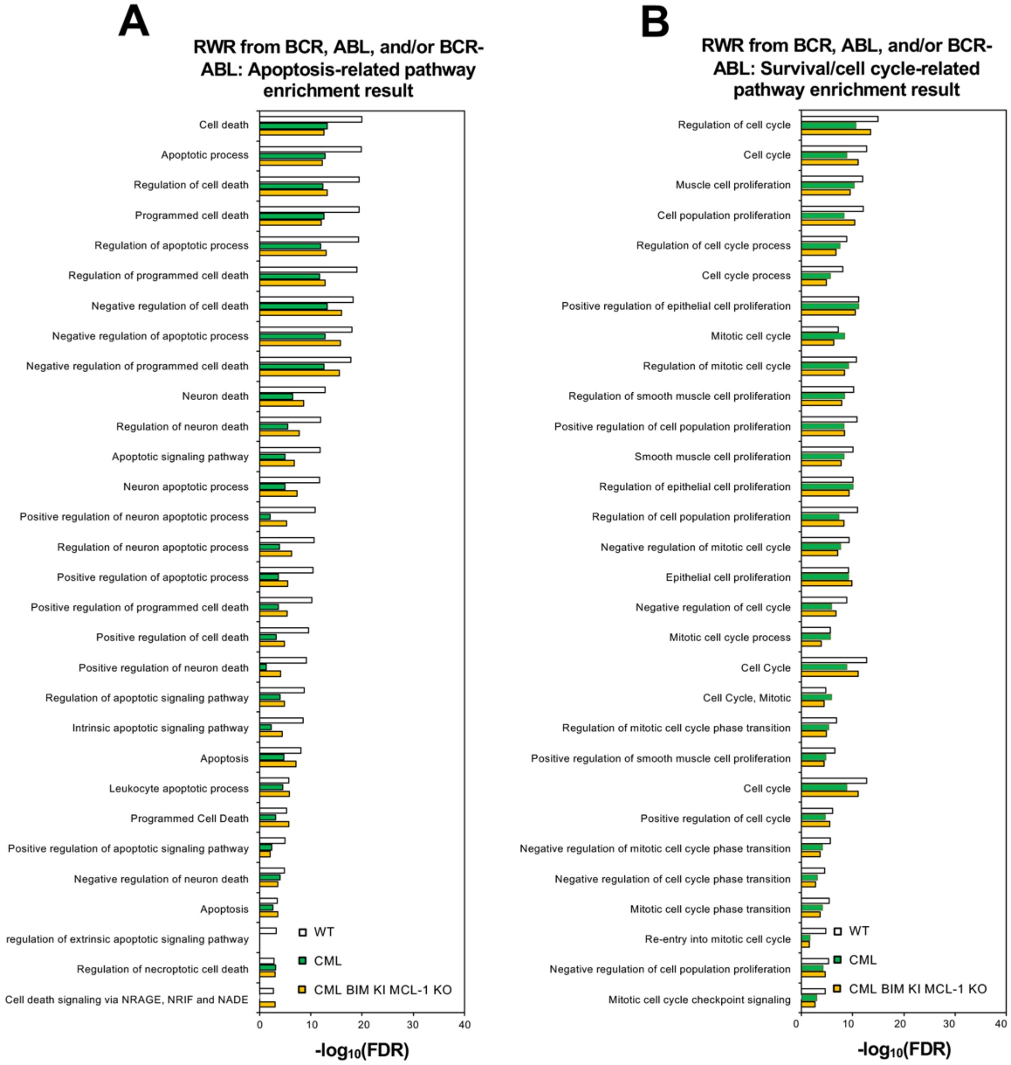


**Figure S3. The effect of BIM KI and MCL-1 KO on BCR/ABL pathway activity in CML.** Bar plots showing the enrichment of (A) apoptosis-related pathways and (B) survival or cell-cycle related pathways based on the RWR score seeded from BCR-ABL and/or BCR and ABL in wild-type (WT; white), CML (green), and CML treated with BIM KI and MCL-1 KO (yellow) networks. CML BIM KI MCL-1 KO demonstrates a mild restoration of apoptosis, survival, and cell cycle-related pathway activities that were lost from WT in the CML network.

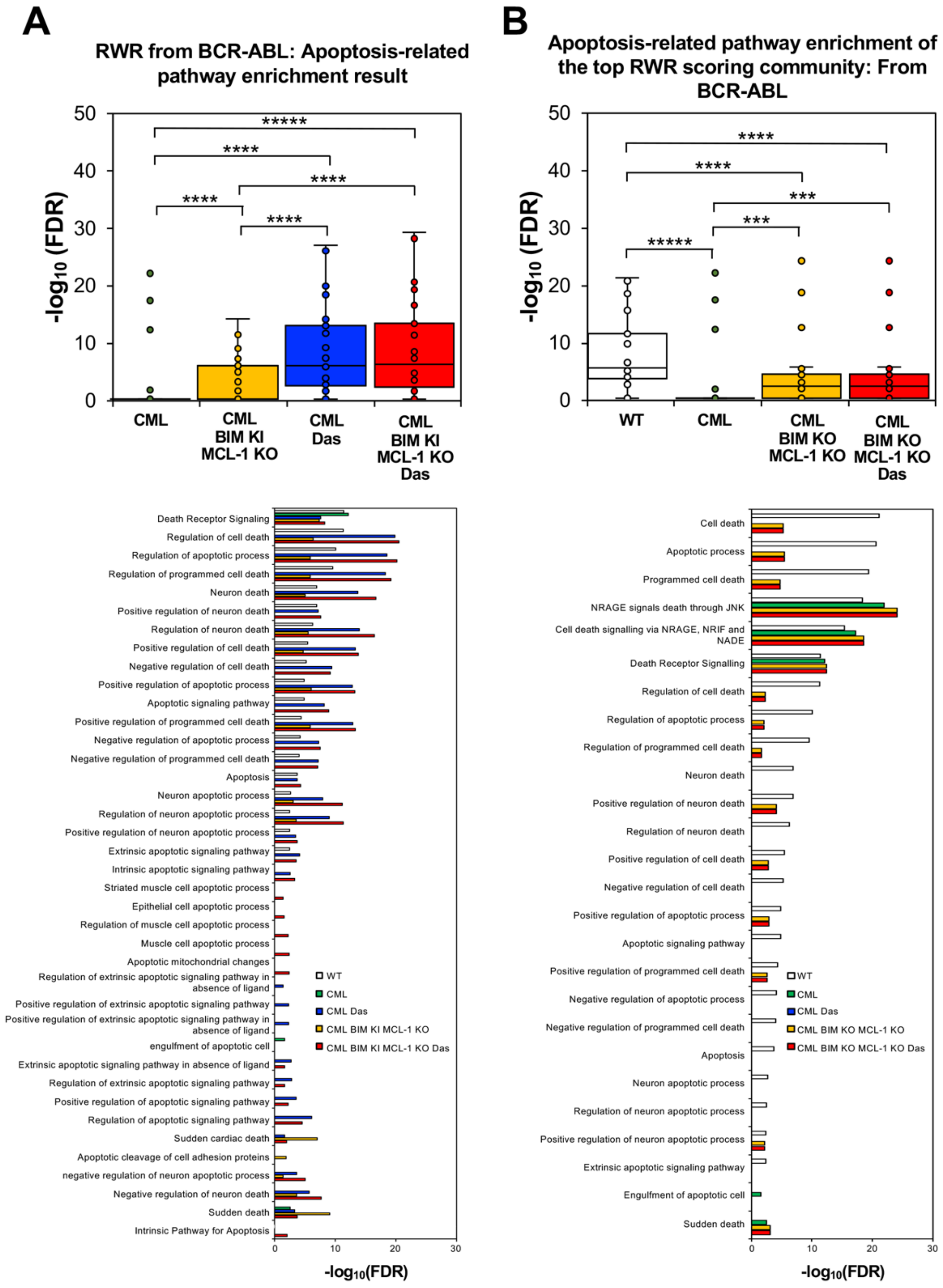


**Figure S4. The effect of targeting BIM and MCL-1 on BCR-ABL signaling based on the community with highest RWR scores.** Summary box plots (top panel) and bar plots (bottom panel) showing the enrichment of apoptosis-related pathways based on the highest RWR scoring community nodes seeded from BCR/ABL and/or BCR and ABL in (A) CML (green), CML treated with BIM KI and MCL-1 KO (yellow), CML treated with Dasatinib (blue), and CML treated with BIM KI and MCL-1 KO and Dasatinib (red) networks and (B) WT (white), CML (green), CML treated with BIM KO MCL-1 KO (yellow) and CML treated with BIM KO MCL-1 KO and Dasatinib (green). CML treated with BIM KI and MCL-1 KO with/without Dasatinib displayed increased apoptotic pathway enrichment compared to CML and other simulated treatments. (*: *p*-value < 0.05; **: *p*-value < 0.01; ***: *p*-value < 0.001; *****: *p*-value < 0.00001)

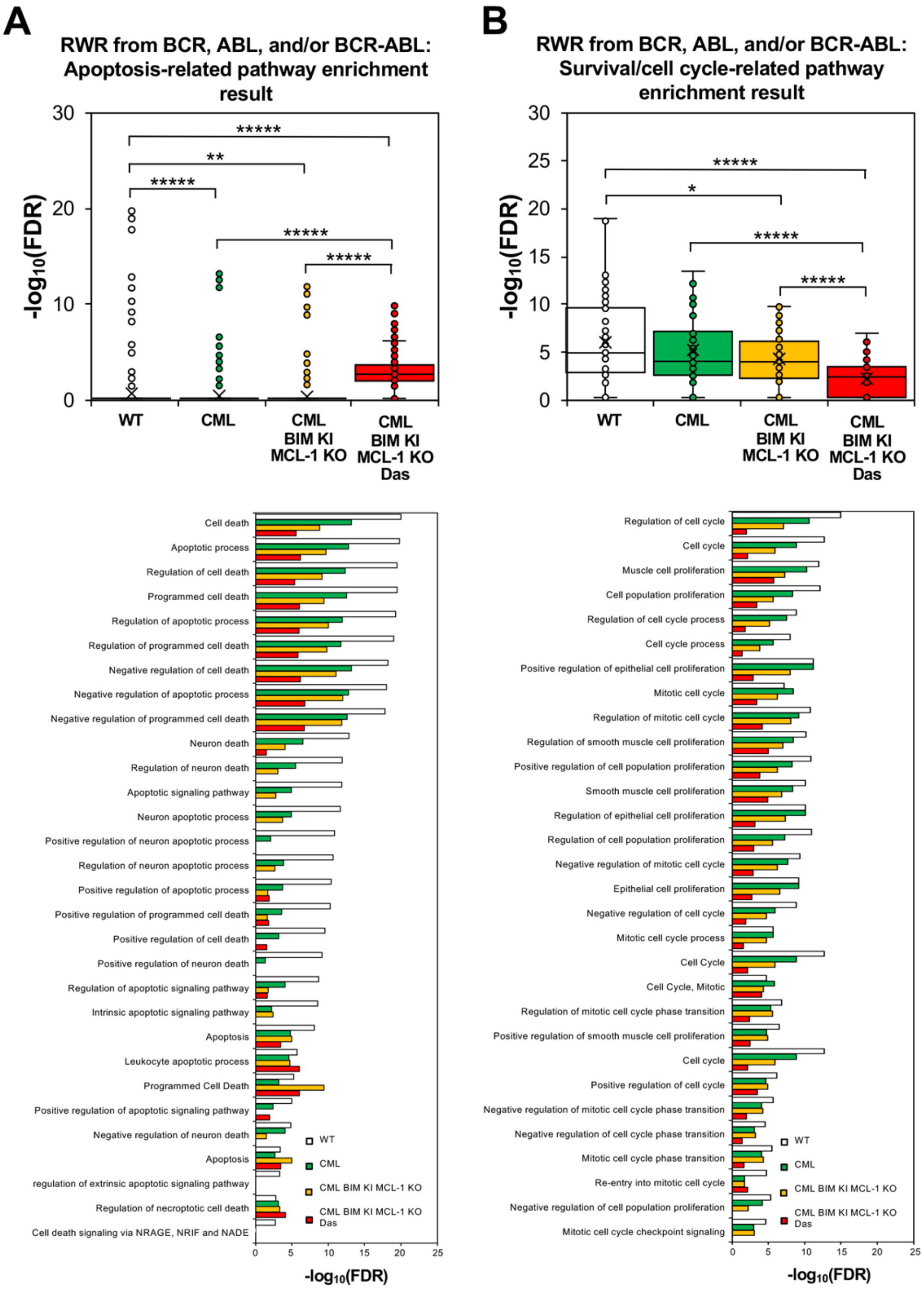


**Figure S5. The effect of BIM KO and MCL-1 KO in CML.** Summary box plots (top panel) and bar plots (bottom panel) showing the enrichment of (A) apoptosis-related pathways and (B) survival or cell-cycle related pathways based on the RWR score seeded from BCR-ABL and/or BCR and ABL in wild-type (WT; white), CML (green), CML treated with BIM KO and MCL-1 KO (yellow) and CML treated with BIM KO and MCL-1 KO with Dasatinib (red) networks. CML BIM KO MCL-1 KO with or without Dasatinib demonstrates a lower level of apoptosis, survival, and cell cycle-related pathway activities than CML or WT networks. (*: $p$-value < 0.05; **: $p$-value < 0.01; ***: $p$-value < 0.001; *****: $p$-value < 0.0001)

**A** **The structure of the simulated heterogeneous multi-layer networks**

| Network | No. of nodes | No. of edges | No. of edges between KEGG and PPI layers |
|---|---|---|---|
| WT | 189 | 141,502 | 14,834 |
| CML | 188 | 136,910 | 14,617 |
| CML BIM KI MCL-1 KO | 189 | 138,085 | 14,615 |
| CML Das | 188 | 135,329 | 14,493 |
| CML BIM KI MCL-1 KO Das | 189 | 136,504 | 14,491 |
| CML BIM KO MCL-1 KO | 188 | 134,354 | 14,438 |
| CML BIM KO MCL-1 KO Das | 187 | 131,777 | 14,256 |

**B**

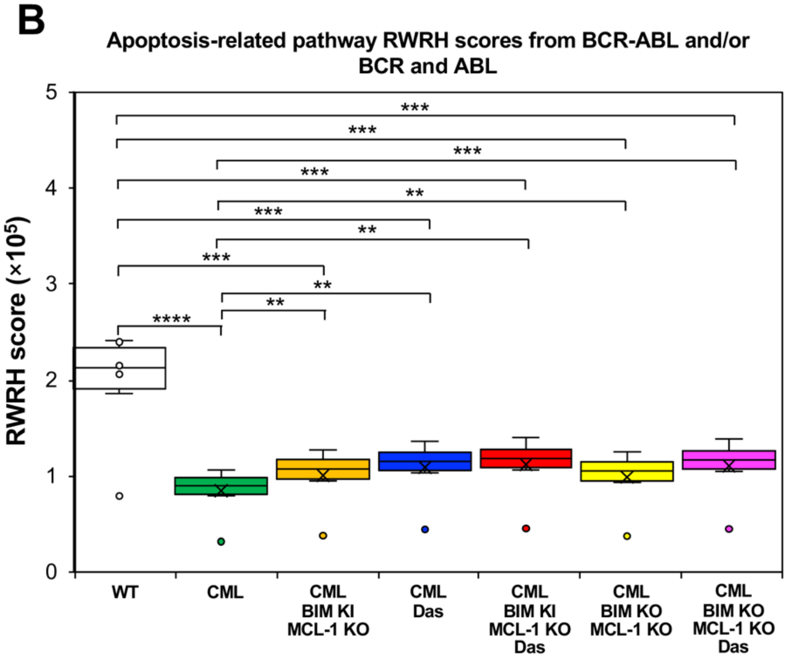


**Figure S6. RWRH z-scores of multi-layer network with KEGG layer generated based on pathway enrichment results.** (A) Table shows the total number of pathway terms in nodes and the number of edges connecting the KEGG pathway terms in the heterogeneous multi-layer network. The number of edges connecting the KEGG layer network to the PPI network is also summarized. (B) Box plot showing the Random-Walk-Restart-Heterogenous (RWRH) z-scores of all apoptosis-related pathways at the KEGG layer network, seeded from BCR-ABL and/or BCR and ABL at the PPI layer. Apoptosis-related pathways are shown to be significantly different between WT network and all CML networks included those under treatment. Apoptosis-related pathways also scored significantly lower to CML networks under any treatment. (**: *p*-value < 0.01; ***: *p*-value < 0.001; ****: *p*-value < 0.0001)

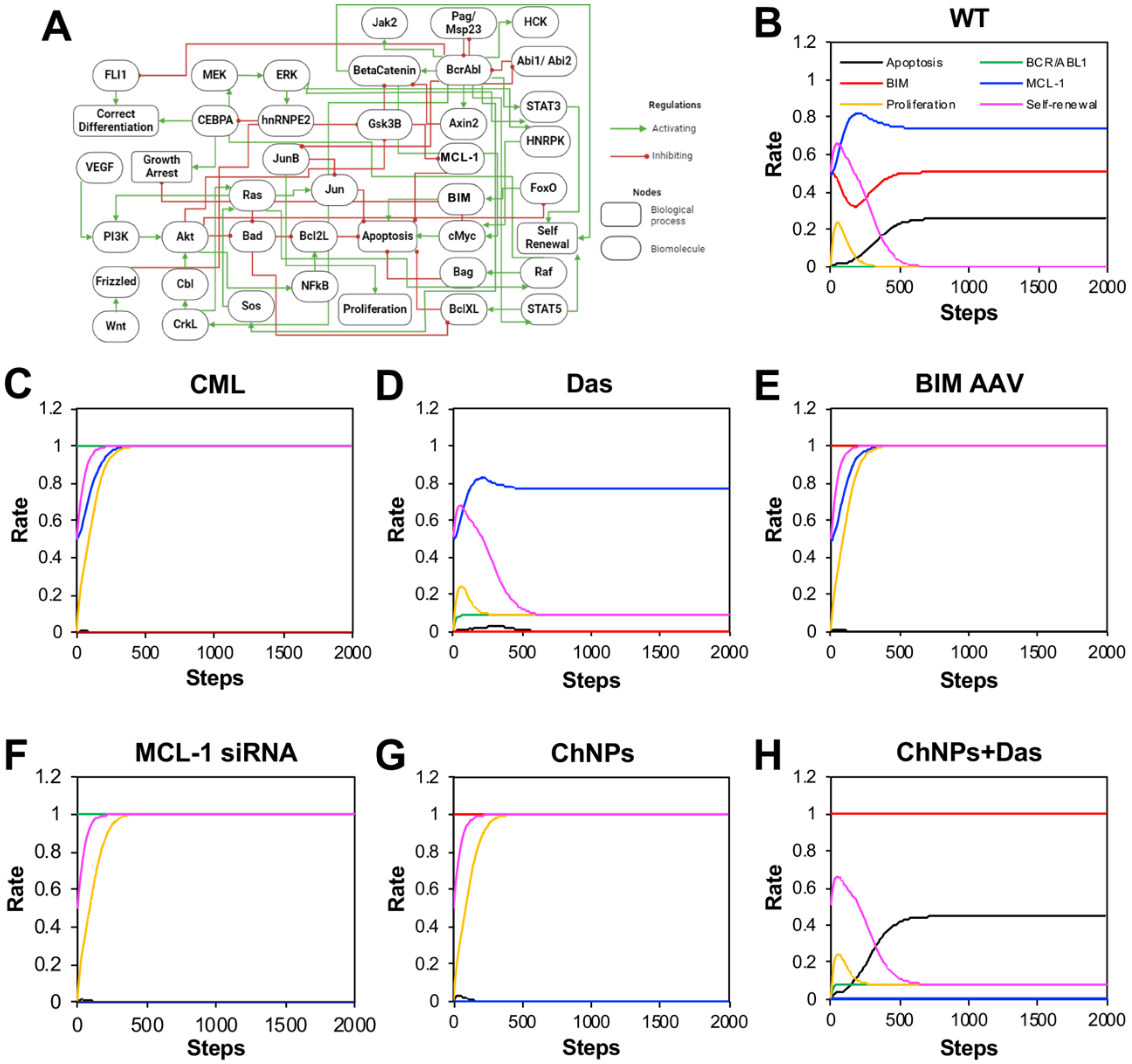


**Figure S7. CML Boolean network simulation.** (A) The final reduced CML Boolean network topology. A green line with an arrowhead represents positive regulation, whereas a red line with a diamond indicates negative regulation. Activation frequencies for three pathway nodes, including Proliferation, Apoptosis, and Self-Renewal, together with three biomolecule nodes, including BCR-ABL, MCL-1, and BIM, were observed in simulated (B) WT, (C) CML, (D) CML treated with Dasatinib, (E) CML treated with BIM AAV, (F) CML treated with MCL-1 siRNA, (G) CML treated with BIM KI MCL-1 KO ChNPs, and (H) CML treated with BIM KI MCL-1 KO ChNPs and Dasatinib conditions.

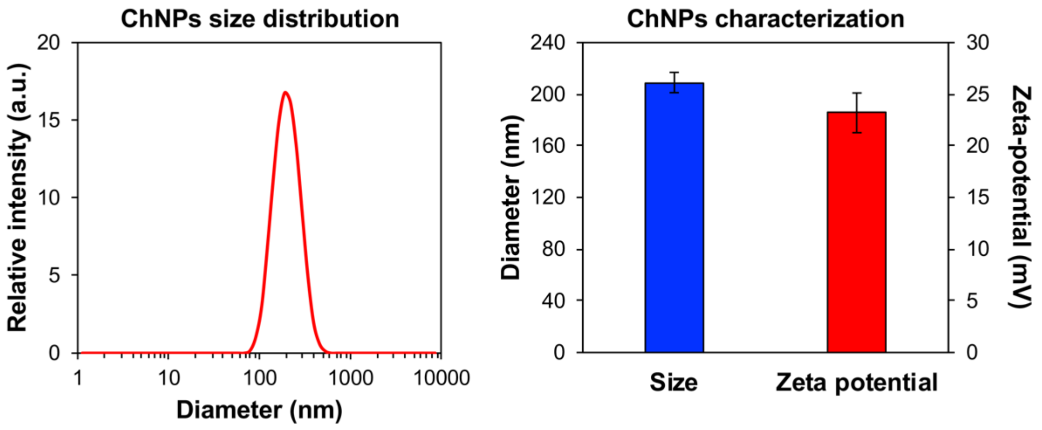


**Figure S8. ChNP size and zeta potential characterization.** ChNPs had a uniform size around 200nm in diameter as observed by the normal curve. Their zeta-potential was positive, as expected due to positive nature of the polymer shell being created.

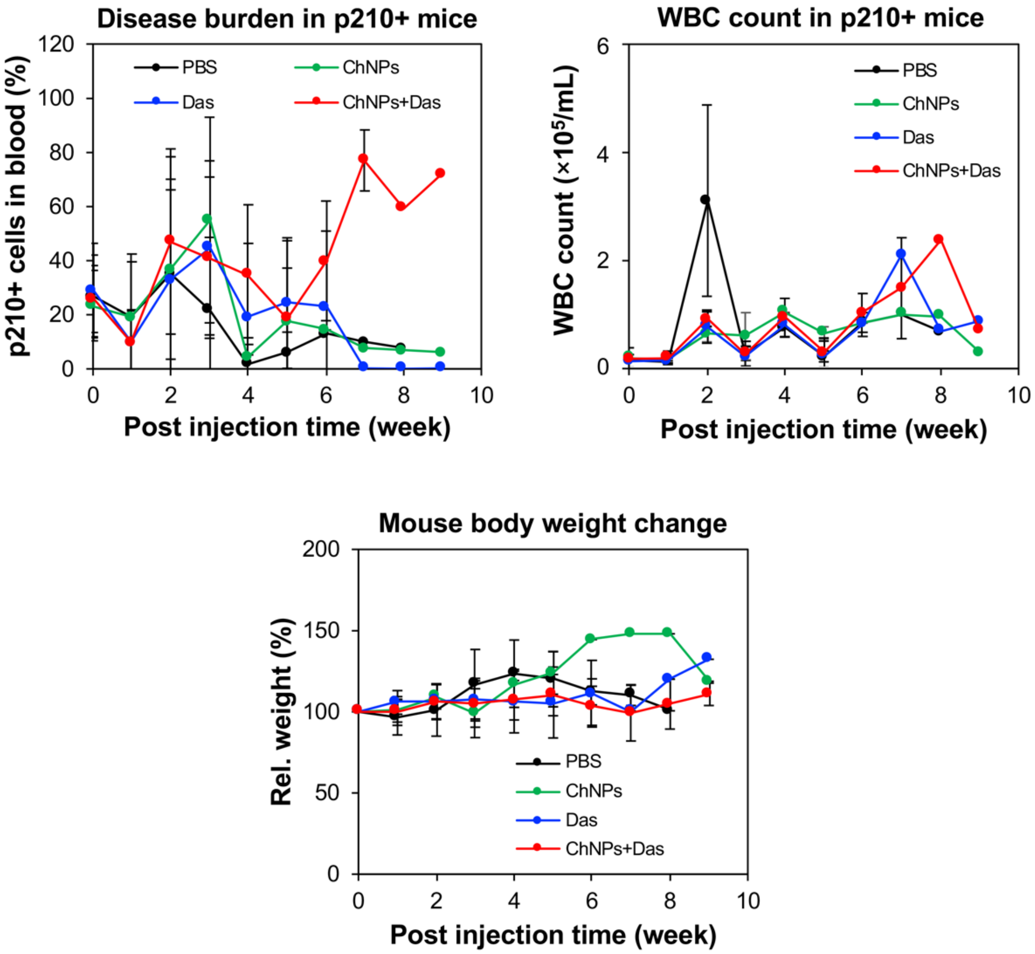


**Figure S9. Disease burden, white blood cell (WBC) counts, and weights of diseased mice compared with weeks after treatment in mice with chronic phase BCR-ABL+ leukemia.** Disease burden appeared higher over time in mice being treated with ChNPs + Das as mice with heavier disease burden were able to survive longer after treatment was removed, while other groups had reduced burden through losing mice in the group. All mice had similar WBC counts expect for PBS mice as they were going through blast crisis. Mice being treated with ChNPs were able to gain the most weight, but weight was largely steady for all groups.

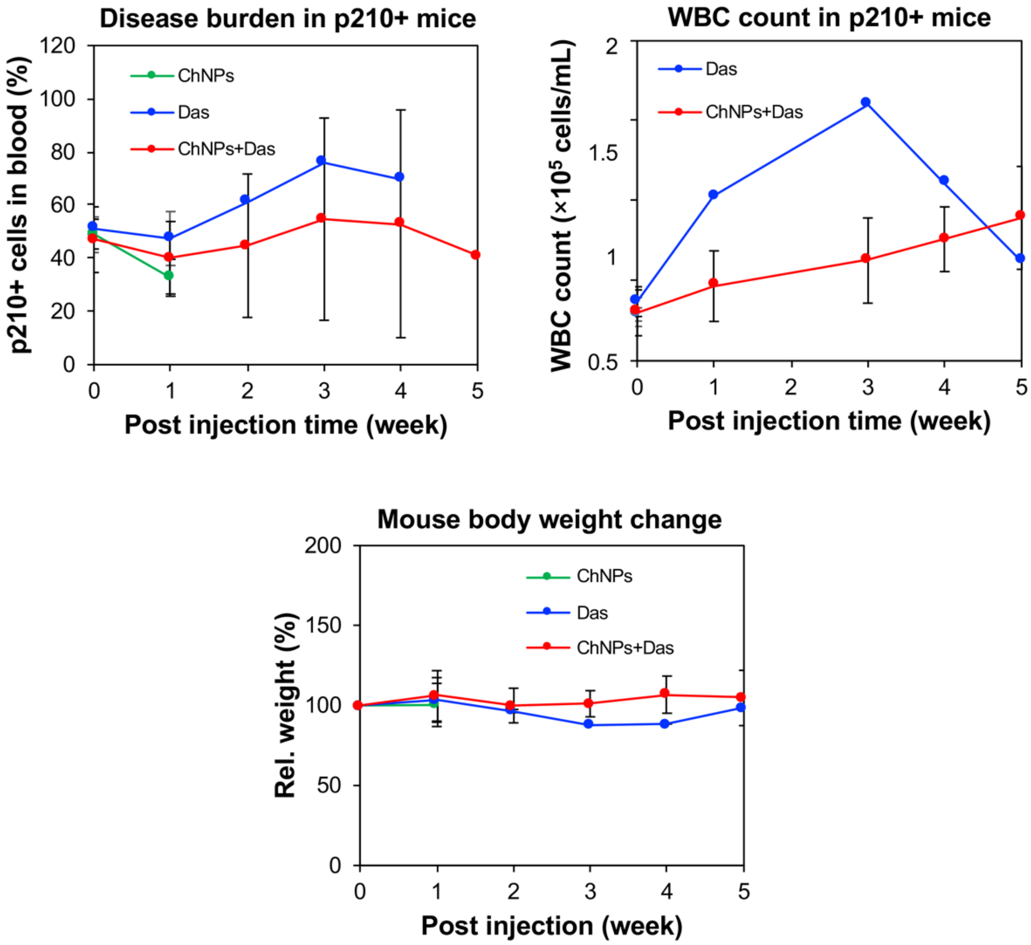


**Figure S10. Disease burden, white blood cell (WBC) counts, and weights of diseased mice compared with weeks after treatment in mice with acute phase BCR-ABL+ leukemia.** Disease burden was highest in the two dasatinib groups as mice with higher disease burden were able to live longer. WBC counts were higher in the dasatinib group compared to the ChNPs + Das group, indicating spreading disease. Body weight was consistent across treatment groups.

**Table S1.** Two-tailed student's *t*-test statistics for p210 (**Fig. 3A iii**)

| | HEPES | Null AAV | Scr siRNA | Null AAV + Scr siRNA | BIM AAV | MCL-1 siRNA | BIM AAV + MCL-1 siRNA | Null/Scr ChNPs | Null/MCL-1 ChNPs | BIM/Scr ChNPs |
|---|---|---|---|---|---|---|---|---|---|---|
| Null AAV | 0.15304 | | | | | | | | | |
| Scr siRNA | 0.26422 | 0.78157 | | | | | | | | |
| Null AAV + Scr siRNA | 0.41491 | 0.12814 | 0.30509 | | | | | | | |
| BIM AAV | 0.47687 | 0.46278 | 0.54533 | 0.7827943 | | | | | | |
| MCL-1 siRNA | 0.13521 | 0.59353 | 0.6375 | 0.6794987 | 0.83408 | | | | | |
| BIM AAV + MCL-1 siRNA | 0.18044 | 0.25179 | 0.38444 | 0.8339913 | 0.68772 | 0.34906 | | | | |
| Null/Scr ChNPs | 0.08207 | 0.33305 | 0.53092 | 0.1649166 | 0.64367 | 0.78735 | 0.20803845 | | | |
| Null/MCL-1 ChNPs | 0.2428 | 0.85204 | 0.93093 | 0.4712769 | 0.61773 | 0.55285 | 0.41485507 | 0.61409 | | |
| BIM/Scr ChNPs | 0.33513 | 0.61254 | 0.61513 | 0.7862966 | 0.91305 | 0.97045 | 0.65616349 | 0.81241 | 0.30605 | |
| BIM/MCL-1 ChNPs | 0.00017 | 0.03982 | 0.09016 | 0.0216902 | 0.0572 | 0.01738 | 0.01091418 | 0.01091 | 0.08295 | 0.01602 |

**Table S2.** Two-tailed student's *t*-test statistics for MiG (**Fig. 3A iv**)

| | 0 nM Das | 5 nM Das | 10 nM Das |
|---|---|---|---|
| 0 nM Das + ChNPs | 0.96237 | | |
| 5 nM Das + ChNPs | | 0.48088 | |
| 10 nM Das + ChNPs | | | 0.630422 |

**Table S3.** Two-tailed student's *t*-test statistics for p210 (**Fig. 3A iv**)

| | 0 nM Das | 5 nM Das | 10 nM Das |
|---|---|---|---|
| 0 nM Das + ChNPs | 0.00878 | | |
| 5 nM Das + ChNPs | | 0.03002 | |
| 10 nM Das + ChNPs | | | 0.019438 |

$p < 0.05$ $p < 0.05$ $p < 0.005$ $p < 0.001$

**Table S4.** Two-tailed student's *t*-test statistics for Ph+ CML (**Fig. 3B i**)

| | HEPES | Null AAV | Scr siRNA | Null AAV + Scr siRNA | BIM AAV | MCL-1 siRNA | BIM AAV + MCL-1 siRNA | Null/Scr ChNPs | Null/MCL-1 ChNPs | BIM/Scr ChNPs | BIM/MCL-1 ChNPs | BIM/MCL-1 ChNPs + Das |
|---|---|---|---|---|---|---|---|---|---|---|---|---|
| Null AAV | 0.37286 | | | | | | | | | | | |
| Scr siRNA | 0.15807 | 0.93588 | | | | | | | | | | |
| Null AAV + Scr siRNA | 0.10385 | 0.15701 | 0.09587 | | | | | | | | | |
| BIM AAV | 0.27745 | 0.28633 | 0.12408 | 0.50568 | | | | | | | | |
| MCL-1 siRNA | 0.08261 | 0.17016 | 0.04611 | 0.33219 | 0.84265 | | | | | | | |
| BIM AAV + MCL-1 siRNA | 0.5378 | 0.88647 | 0.91632 | 0.21665 | 0.36719 | 0.26035 | | | | | | |
| Null/Scr ChNPs | 0.6151 | 0.19566 | 0.29605 | 0.43927 | 0.62491 | 0.53993 | 0.23458 | | | | | |
| Null/MCL-1 ChNPs | 0.01004 | 0.00315 | 0.00954 | 0.02373 | 0.02266 | 0.01555 | 0.00672 | 0.00084 | | | | |
| BIM/Scr ChNPs | 0.00672 | 0.00645 | 0.00427 | 0.02202 | 0.01851 | 0.01145 | 0.02023 | 0.00438 | 0.05302 | | | |
| BIM/MCL-1 ChNPs | 0.00815 | 0.01112 | 0.0042 | 0.01882 | 0.00952 | 0.00874 | 0.02253 | 0.00384 | 0.76997 | 0.03731 | | |
| BIM/MCL-1 ChNPs + Das | 0.00974 | 0.00405 | 0.00861 | 0.02184 | 0.01886 | 0.01406 | 0.0067 | 0.00016 | 0.01279 | 0.02405 | 0.23993 | |
| Das | 0.05595 | 0.06286 | 0.04788 | 0.06372 | 0.03656 | 0.04247 | 0.08497 | 0.01202 | 0.09677 | 0.31277 | 0.04303 | 0.04051 |

**Table S5.** Two-tailed student's *t*-test statistics for Ph+ CML BC (**Fig. 3B ii**)

| | HEPES | Null AAV | Scr siRNA | Null AAV + Scr siRNA | BIM AAV | MCL-1 siRNA | BIM AAV + MCL-1 siRNA | Null/Scr ChNPs | Null/MCL-1 ChNPs | BIM/Scr ChNPs | BIM/MCL-1 ChNPs | BIM/MCL-1 ChNPs + Das |
|---|---|---|---|---|---|---|---|---|---|---|---|---|
| Null AAV | 0.79527 | | | | | | | | | | | |
| Scr siRNA | 0.82388 | 0.7287 | | | | | | | | | | |
| Null AAV + Scr siRNA | 0.65612 | 0.99925 | 0.4933 | | | | | | | | | |
| BIM AAV | 0.60398 | 0.41459 | 0.55168 | 0.3622267 | | | | | | | | |
| MCL-1 siRNA | 0.47782 | 0.58427 | 0.62743 | 0.2472423 | 0.63761 | | | | | | | |
| BIM AAV + MCL-1 siRNA | 0.07064 | 0.87231 | 0.40723 | 0.8019707 | 0.4788 | 0.21193 | | | | | | |
| Null/Scr ChNPs | 0.73931 | 0.89761 | 0.37586 | 0.68027 | 0.39347 | 0.16377 | 0.66896238 | | | | | |
| Null/MCL-1 ChNPs | 0.06489 | 0.45028 | 0.06111 | 0.1385819 | 0.25522 | 0.00194 | 0.28443014 | 0.05935 | | | | |
| BIM/Scr ChNPs | 0.05142 | 0.48601 | 0.0279 | 0.2005163 | 0.25422 | 0.00355 | 0.26272418 | 0.07211 | 0.79819 | | | |
| BIM/MCL-1 ChNPs | 0.02169 | 0.06661 | 0.00585 | 0.0072109 | 0.05986 | 0.00295 | 0.04527416 | 0.000034 | 0.00928 | 0.01285 | | |
| BIM/MCL-1 ChNPs + Das | 0.016 | 0.05482 | 0.00661 | 0.0049333 | 0.06007 | 0.00168 | 0.03348313 | 0.00056 | 0.00429 | 0.00891 | 0.10019 | |
| Das | 0.07694 | 0.43028 | 0.00389 | 0.162502 | 0.20744 | 0.01113 | 0.26912279 | 0.04159 | 0.45116 | 0.31066 | 0.01163 | 0.01096 |

**Table S6.** Two-tailed student's *t*-test statistics for Ph- AML (**Fig. 3B iii**)

| | HEPES | Null AAV | Scr siRNA | Null AAV + Scr siRNA | BIM AAV | MCL-1 siRNA | BIM AAV + MCL-1 siRNA | Null/Scr ChNPs | Null/MCL-1 ChNPs | BIM/Scr ChNPs | BIM/MCL-1 ChNPs | BIM/MCL-1 ChNPs + Das |
|---|---|---|---|---|---|---|---|---|---|---|---|---|
| Null AAV | 0.21671 | | | | | | | | | | | |
| Scr siRNA | 0.22706 | 0.23874 | | | | | | | | | | |
| Null AAV + Scr siRNA | 0.52562 | 0.99164 | 0.05224 | | | | | | | | | |
| BIM AAV | 0.29412 | 0.97032 | 0.28838 | 0.9786762 | | | | | | | | |
| MCL-1 siRNA | 0.20011 | 0.0057 | 0.0545 | 0.1559211 | 0.07009 | | | | | | | |
| BIM AAV + MCL-1 siRNA | 0.15434 | 0.30651 | 0.60145 | 0.4132965 | 0.22036 | 0.12634 | | | | | | |
| Null/Scr ChNPs | 0.32521 | 0.40916 | 0.72848 | 0.1589389 | 0.39188 | 0.13706 | 0.71450082 | | | | | |
| Null/MCL-1 ChNPs | 0.0217 | 0.01028 | 0.00366 | 0.0056175 | 0.00615 | 0.00922 | 0.07455442 | 0.00681 | | | | |
| BIM/Scr ChNPs | 0.22871 | 0.30808 | 0.98984 | 0.2223006 | 0.46945 | 0.05581 | 0.66451286 | 0.84273 | 0.0235 | | | |
| BIM/MCL-1 ChNPs | 0.00718 | 0.00198 | 0.00121 | 0.0028702 | 0.00113 | 0.00243 | 0.02961444 | 0.0068 | 0.01034 | 0.00804 | | |
| BIM/MCL-1 ChNPs + Das | 0.0064 | 0.00152 | 0.00114 | 0.0027555 | 0.00103 | 0.00199 | 0.02902852 | 0.0072 | 0.01273 | 0.00714 | 0.11533 | |
| Das | 0.06115 | 0.03877 | 0.08243 | 0.0367278 | 0.0944 | 0.0145 | 0.71335674 | 0.26132 | 0.04554 | 0.00356 | 0.01139 | 0.00986 |

$p < 0.05$ $p < 0.05$ $p < 0.005$ $p < 0.001$

**Table S7.** Two-tailed student's t-test statistics for chronic leukemia survival data (**Fig. 4B**)

| | PBS | ChNPs | Das | ChNPs + Das |
|---|---|---|---|---|
| ChNPs | 0.0758 | | | |
| Das | 0.0014 | 0.0025 | | |
| ChNPs + Das | 0.0055 | 0.0409 | 0.0025 | |

**Table S8.** Two-tailed student's *t*-test statistics for acute leukemia survival data (**Fig. 4B**)

| | PBS | ChNPs | Das | ChNPs + Das |
|---|---|---|---|---|
| ChNPs | 0.00972 | | | |
| Das | 0.16708 | 0.38195 | | |
| ChNPs + Das | 0.04536 | 0.08262 | 0.27184 | |

**Table S9.** Two-tailed student's *t*-test statistics for pathology study (Liver, Day 11, **Fig. 4D**)

| | Healthy | PBS | ChNPs | ChNPs + Das |
|---|---|---|---|---|
| PBS | * (0.0184) | | | |
| ChNPs | NS | NS | | |
| ChNPs + Das | NS | * (0.0123) | NS | |
| Das | NS | * (0.0103) | NS | NS |

**Table S10.** Two-tailed student's t-test statistics for pathology study (Liver, Day 47, **Fig. 4D**)

| | Healthy | ChNPs + Das |
|---|---|---|
| ChNPs + Das | NS | |
| Das | ** (0.0019) | ** (0.0056) |

**Table S11.** Two-tailed student's *t*-test statistics for pathology study (Spleen, Day 11, **Fig. 4D**)

| | Healthy | PBS | ChNPs | ChNP + Das |
|---|---|---|---|---|
| PBS | *** (0.001) | | | |
| ChNPs | ** (0.03) | NS | | |
| ChNPs + Das | NS | *** (0.001) | *** (0.001) | |
| Das | NS | ** (0.03) | ** (0.012) | NS |

**Table S12.** Two-tailed student's *t*-test statistics for Pathology study (Spleen, Day 47, **Fig. 4D**)

| | Healthy | ChNPs + Das |
|---|---|---|
| ChNPs + Das | NS | |
| Das | * (0.0253) | * (0.047) |

$p < 0.05$  $p < 0.05$  $p < 0.005$  $p < 0.001$
